# Mesenchymal Stromal Cell Therapeutic Potential to Restore Neurovascular Integrity in Machado-Joseph Disease

**DOI:** 10.64898/2026.01.31.703043

**Authors:** Inês Barros, Daniela Gonzaga, Diana Lobo, Sara Lopes, Inês Martins, António Silva, Carina Henriques Silva, Célia Gomes, Rui Jorge Nobre, Luí Pereira de Almeida, Catarina Oliveira Miranda

**Affiliations:** CNC-UC - Center for Neuroscience and Cell Biology, University of Coimbra, Portugal; GeneT – Gene Therapy Center of Excellence, Portugal; CiBB - Center for Innovative Biomedicine and Biotechnology, Coimbra, Portugal; iiiUC - Institute for Interdisciplinary Research, University of Coimbra, Portugal; PDBEB - University of Coimbra, Institute for Interdisciplinary Research, Doctoral Programme in Experimental Biology and Biomedicine, Portugal; Faculty of Pharmacy, University of Coimbra, Portugal; iCBR - Coimbra Institute for Clinical and Biomedical Research; ViraVector - Viral Vector for Gene Transfer Core facility, University of Coimbra, Portugal

**Keywords:** Blood-brain barrier (BBB), Machado-Joseph disease (MJD), Mesenchymal Stromal Cells (MSC), Neurovascular dysregulations, Polyglutamine disorders (PolyQ)

## Abstract

**Background:** Machado-Joseph disease (MJD), or Spinocerebellar Ataxia type 3 (SCA3), is a neurodegenerative disorder caused by an expansion of the CAG repeat in the *MJD1/ATXN3* gene, which encodes for a polyglutamine-expansion in Ataxin-3. Similar to what occurs in other neurodegenerative disorders, the cerebrovascular system, particularly the blood-brain barrier (BBB), is impaired in MJD. BBB is an important cellular barrier that controls homeostasis of the central nervous system, and its disruption compromises neuronal function. To date, there is no disease-modifying therapy for MJD. Our group previously demonstrated that MSC administration can ameliorate the phenotype of MJD transgenic mice. However, the effect of MSC on the neurovascular unit and BBB integrity remains unexplored in the context of MJD. In the present work, we aimed to evaluate the therapeutic potential of MSC in mitigating vascular abnormalities and BBB disruption in MJD.

**Methods:** Eight-month-old MJD transgenic mice were treated with two consecutive bone marrow-MSC intravenous injections (1 week apart). Behavioral tests to assess motor coordination were conducted before and after treatment. Vascular structure, function, and BBB disruption were evaluated by immunohistochemistry and western blot.

**Results:** MSC administration partially restored vascular impairment in MJD transgenic mice. Treated mice exhibited improved gait and motor performance footprint and beam walking tests) and concurrently attenuated vascular dysfunction. Specifically, MSC-treated mice exhibited a decrease in collagen IV surface area and stabilized cerebellar blood supply. Additionally, MSC treatment diminished BBB permeability to the Evans blue (EB) dye in MJD mice and restored the levels of important adherent and tight junction-associated proteins in the cellular membrane, including vascular endothelial cadherin (VE-cadherin), claudin-5, and occludin, in a sex-specific manner.

**Conclusions:** our results highlight significant cerebrovascular dysregulations in MJD, which are sex-dependent and progress with the disease stage. Moreover, our findings suggest that MSC administration partially reverted vascular impairments, providing valuable insights into the mechanisms underlying the therapeutic potential of MSC in MJD.

**Graphical Abstract:** MSC treatment partially reverts neurovascular dysregulation in MJD. MJD mice exhibit increased vessel coverage, increased BBB permeability, and altered TJ and AJ expression and subcellular localization. MSC administration modulates vessel coverage, reduces BBB leakage, and partially normalizes TJ/AJ subcellular localization.

<u>List of abbreviations:</u> AJ – Adherent junction; BBB – Blood-Brain barrier; MJD- Machado-Joseph disease; MSC – mesenchymal stromal cell; TJ – thigh junction. Created with BioRender.com

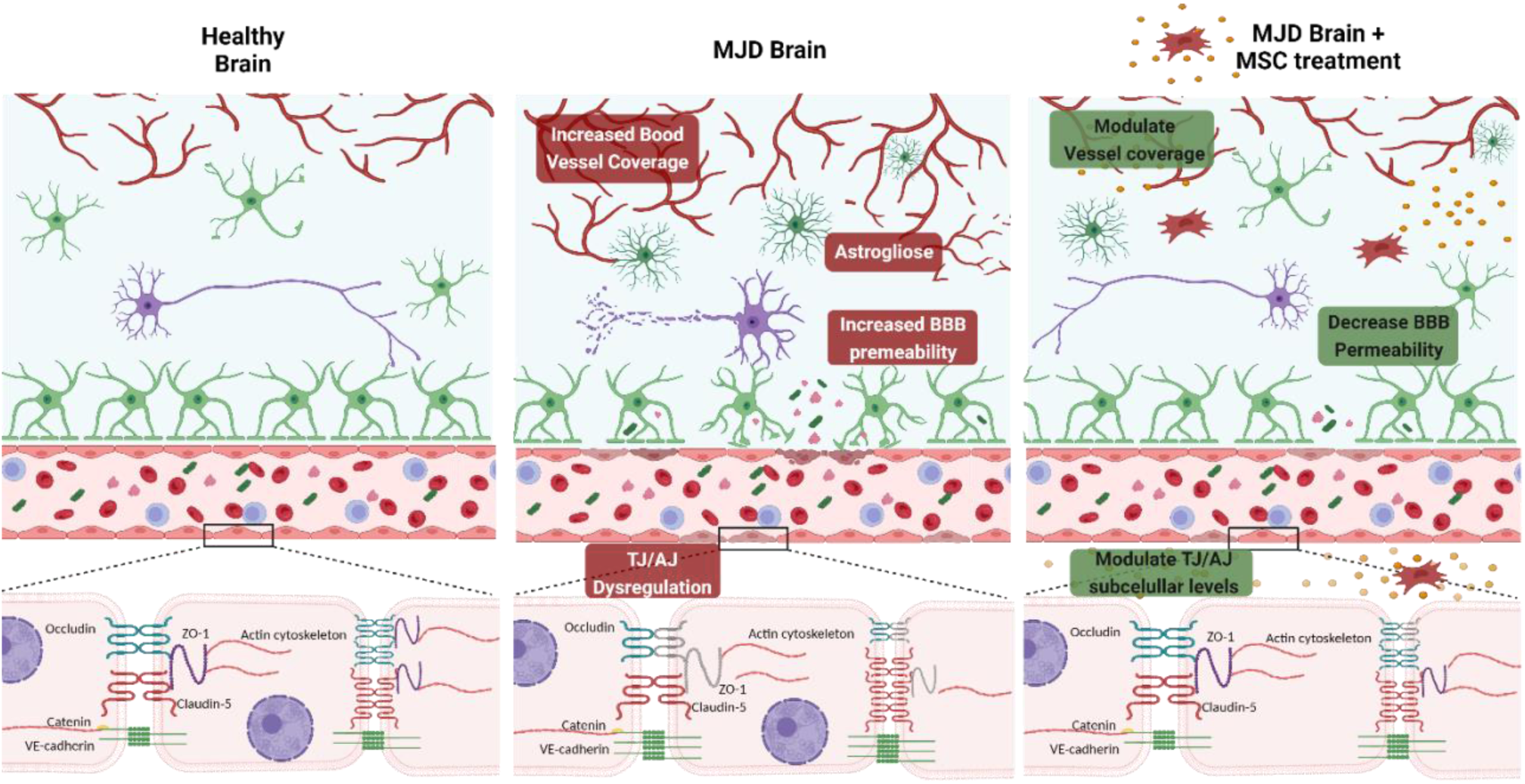

## Introduction

Machado-Joseph disease/spinocerebellar ataxia type 3 (MJD/SCA3), is the most common autosomal dominantly-inherited ataxia worldwide. It is caused by an unstable expansion of the CAG repeat in the *MJD1/ATXN3* gene, encoding an abnormally long polyglutamine (PolyQ) stretch in ataxin-3 (1, 2). Usually, patients start exhibiting the first symptoms in young or mid-adulthood, and the disease slowly progresses over the following years (21 years on average) (3). Some of the most common clinical features are progressive ataxia, dysarthria, dysphagia, and atrophy, among other motor and non-motor symptoms (4, 5). While the exact mechanism underlying mutant ataxin-3 toxicity remains largely unknown, it is well established that the presence of mutant protein triggers the formation of aggregates and disrupts crucial cellular mechanisms, culminating in widespread neuronal degeneration (reviewed in (6)). Indeed, several brain regions are affected in MJD, including the cerebellum, brainstem, substantia nigra, thalamus, striatum, and spinal cord (Sudarsky and Coutinho, 1995, Durr et al., 1996, Alves et al., 2008, Koeppen, 2018).

Increasing evidence indicates that vascular dysfunctions affecting the neurovascular unit (NVU) and, specifically, the blood-brain barrier (BBB) are implicated in the pathology of several neurogenerative disorders. BBB is a selective cellular barrier that restricts the passage of ions, molecules, and cells from the bloodstream to the brain. It is formed by a layer of endothelial cells with no fenestrations between them due to the presence of tight junctions (TJ) and adherent junctions (AJ). Endothelial cells are supported by pericytes, and both are embedded by the basement membrane and astrocyte end-feet. Additionally, perivascular neurons and microglial cells can also influence BBB dynamics. BBB disruption leads to the accumulation of toxic blood-borne proteins in the brain’s parenchyma and increases its exposure to viruses, bacteria, and peripheral immune cells, ultimately contributing to neuronal injury, neurotoxicity, and neuroinflammation (reviewed in (7)). In a previous study, our group found evidence of BBB dysregulation in both late-stage MJD transgenic (Tg) mice (Tg-ATXN3-69Q model (8)), and post-mortem brain samples from MJD patients (9). Moreover, cerebral blood flow was shown to be impaired in patients with spinocerebellar ataxias, including MJD (10–12). Nevertheless, a deeper understanding of the neurovascular alterations in MJD, at a structural and functional level, and how they evolve with the disease progression remains imperative.

Despite all research efforts to date, no therapy can cure or efficiently delay MJD progression (13). Nevertheless, over the last decades, mesenchymal stromal cells (MSC) have been highlighted as promising therapeutic agents for the treatment of several neurodegenerative diseases, including PolyQ disorders. MSC are multipotent cells that promote tissue regeneration, repair, and turnover through their paracrine activity by increasing cell survival, inducing neurogenesis, and modulating the immune system in these disorders (reviewed in (14)). Further supporting their therapeutic potential, both preclinical (15–17) and clinical studies (18–21) have demonstrated that MSC are safe and may even delay the progression of MJD and other SCAs. Still, MSC’s beneficial effects seem to be transient, as a considerable number of patients were reported to regress to their status before transplantation (18). Accordingly, we have previously shown that MSC transplantation can promote sustained amelioration of MJD symptoms in MJD Tg-ATXN3-69Q mice only when administered repeatedly, possibly because of the short lifetime of MSCs after in vivo transplantation (15). In line with these results, Li et al. demonstrated that repeated pre-symptomatic transplantations of umbilical cord-derived MSC in Tg homozygous MJD mice (YAC-MJD84.2 (22)) led to prolonged alleviation of MJD symptoms (16). However, to improve the efficacy of MSC-based therapy, it is necessary to further explore and identify the mechanisms by which MSC alleviate MJD.

Several studies have demonstrated that MSC have the potential to rescue BBB in acute and chronic neurodegenerative disorders such as ischemic stroke, traumatic brain injury, multiple sclerosis, Parkinson’s disease (PD), and Alzheimer’s disease (AD) (reviewed in (23)). Nevertheless, MSC’s therapeutic effect on vascular and BBB disruption in MJD has not yet been investigated. Thus, in this study we aimed to not only deepen our understanding of cerebrovascular dysregulation in MJD, but also evaluate whether MSC could alleviate such vascular dysregulations, elucidating on the putative mechanisms underlying the MSC therapeutic effects in a MJD mouse model.

Our results reveal significant cerebrovascular changes in MJD a mouse model found, including increased vascular density and alternated vascular function, along with alteration in TJ-and AJ-associated protein expression, subcellular localization and cleavage. Remarkably, many of these neurovascular impairments occurred before major BBB breakdown. Furthermore, our findings demonstrate that repeated administration of MSC not only alleviates MJD motor and gait impairments, but also partially ameliorates vascular abnormalities and restores BBB integrity. Hence, the development of therapies that target vascular dysfunction may potentially alleviate the pathology of MJD and other PolyQ disorders.

## Methods and materials

### Animals

A transgenic animal model (C57BL/6 background) expressing N-terminal truncated human ataxin-3 with 69 CAG repeats (Tg-ATXN3-69Q) preceded by the hemagglutinin (HA) epitope was used in this study (8). The transgene expression is driven by the L7 promoter, specific for cerebellar Purkinje cells. A colony of this model was maintained in the animal facility in the Centre for Neuroscience and Cell Biology (CNC) of the University of Coimbra through backcrossing heterozygous males with C57BL/6 females. For this animal model, the genotype was confirmed by PCR (Polymerase Chain Reaction) analysis using a Veriti™ 96-Well Fast Thermal Cycler (Thermo Fisher Scientific). Electrophoresis was performed on 1% agarose gel using GreenSafe Premium (NZYTech). Bands were visualized in a Gel Doc™ EZ System (Bio-Rad).

Animals were housed in temperature- and humidity-controlled rooms (22 ± 2 °C, 55 ± 15%) and maintained under a 12 h Light/12 h dark cycle. with food and water provided *ad libitum*. Unless otherwise stated, all groups included age-matched male and female mice.

All animal experimental protocols were carried out following the European Community Council Directive (2010/63/EU) for the care and use of laboratory animals, transposed into Portuguese law in 2013 (Decree Law 113/2013). Additionally, animal procedures were previously approved by the Responsible Organization for Animal Welfare of the Faculty of Medicine and CNC of the University of Coimbra (ORBEA and FMUC/CNC, Coimbra, Portugal; ORBEA_66_2015/22062015 and ORBEA_289_2021/10122021). The researchers received adequate training (FELASA-certified course) and certification to perform the experiments from Portuguese authorities (Direcção Geral de Alimentação e Veterinária, Lisbon, Portugal). All efforts were made to minimize animal suffering.

### In vivo experimental design

For the longitudinal characterization of vascular dysregulations in the Tg-ATXN3-69Q mouse model, Wiled-type (WT) and transgenic (Tg) mice of three different age groups were analyzed: a 2-week-old set (referred as 0.5-month-old mice); a 2.5-month-old set and an 8-month-old set. These were the average ages at which mice were sacrificed and tissues were analyzed post-mortem in each set. To evaluated MSC potential to ameliorate vascular dysregulations a group of animals of 8-month-old set were treated with MSC as described below.

#### Mesenchymal Stromal Cells: Cell culture and preparations

MSC were previously isolated from the bone marrow of 6-to 8-week-old wild-type mice (C57BL/6 background) of both sexes and were characterized to guarantee that the population met the criteria of the International Society for Cellular Therapy (24), as reported previously (15). Cells were expanded in Dulbecco’s Modified Eagle Medium: Nutrient Mixture F12 (Gibco) supplemented with 10% fetal bovine serum (FBS, Invitrogen), 100 U/mL penicillin, 100 μg/mL streptomycin (Invitrogen), 20 ng/mL fibroblast growth factor (FGF, PeproTech), 20 ng/ml epidermal growth factor (PeproTech), and 2% B-27 (Gibco), at 37 °C under a humidified atmosphere containing 5% CO2. All experiments throughout this study used MSC from the same batch in passages between 18 to 22.

#### Intravenous injection of MSC

Post-symptomatic 6–7-month-old Tg-ATXN3-69Q mice were transplanted with 4-6 x 10^7^ MSC/kg resuspended in Hanks’ balanced salt solution (HBSS, Sigma). Mice received two intravenous (IV) injections of MSC with 7-10 days of interval between injections. Mice of two different ages were analyzed. We will refer to treated set as 8-month-old animals, since this was the average age at which mice were sacrificed and tissues were analyzed post-mortem. MSC expressing luciferase were used for the bioluminescence studies. For further details, see the schematic experimental timeline in Figure 1A.

**Figure 1.**
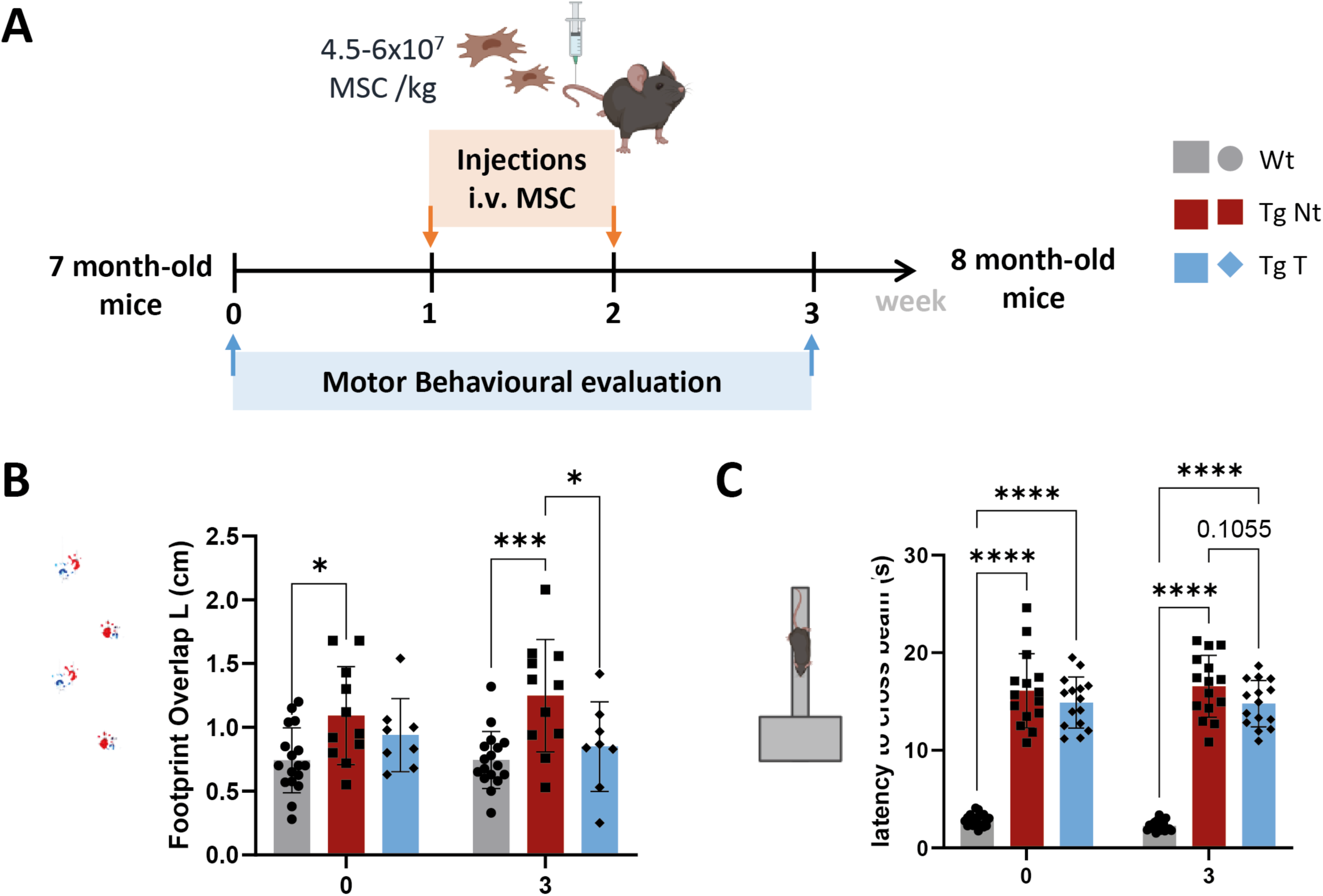
MSC alleviated motor coordination and balance deficits in Tg-ATXN3-69Q Mice. Schematic representation of the experimental procedure (A): Motor performance of 8-month-old Wt, non-treated, and MSC-treated Tg mice were evaluated one week before (week 0) and one week after the last treatment with MSC (week 3). Gait was assessed through footprint left overlap (Wt n=17 vs Tg Nt n=10 vs Tg T n=8) (B). Balance and motor performance as evaluated by beam-walking (Wt n=21 vs Tg Nt n=16 vs Tg T n=15) (C). Data presented as mean ± SD. One-way ANOVA followed by Tukey’s test or pearson correlation. p>0.05 = not significant; * p<0.05, **p< 0.01, ***p< 0.001, ****p< 0.0001.

### Indocyanine Green (ICG) studies: injection and quantification

Mice were intravenously (IV) injected in the caudal vein with a fluorescent contrast agent, indocyanine green (ICG, 7.5 mg/kg) resuspended in water. ICG fluorescence was determined 10, 30, and 60 min after injection using the IVIS Spectrum imaging system (Xenogen, Perkin Elmer, MA, USA). ICG signal was quantified in three different areas of interest: regions of interest (ROI) 1 – entire brain; ROI 2 – cerebellum; and ROI 3 – maximum signal automatically selected by IVIS software. Only the data form ROI 2 is presented in this study. The imaging data acquired from IVIS were quantified in units of radiant efficiency, which means photons per second per square cm per steradian/microwatts per square cm. Animals of the 2.5- and 8-month-old set were analyzed.

### Tissue preparation

Tg-ATXN3-69Q MJD mice with 2.5- and 8-month-old were terminally anesthetized via the intraperitoneal route with an overdose of a mixture of ketamine (Clorketam 1000, Vétoquinol, Lure, France) and xylazine (Rompun, Bayer, Leverkusen, Germany) and transcardially perfused with ice-cold Phosphate-buffered saline solution (PBS) 1x, (pH 7.4). Perfused brains were harvested and cut in halves. The left hemispheres were preserved in tissue-tek (Sakura) at −80 °C to be posteriorly analyzed by immunofluorescence. The right hemispheres were dissected to collect the cerebellum. Cerebellar samples were stored at −80 °C for further protein and ribonucleic acid (RNA) extraction and analysis.

### Protein extraction and Western Blotting

Total protein extracts and subcellular protein fractions were obtained from the right cerebellar hemisphere, as detailed in the Supplemental Methods and Materials section. Bradford protein assay (BioRad) was used to determine protein concentration.

For the Western Blotting analysis, 40-50 micrograms of total protein extracts, 30-40 micrograms of cytoplasmic and membrane protein extracts, and 15-20 micrograms of cytoskeletal protein extracts were loaded per lane and resolved by electrophoresis on an SDS-polyacrylamide gel (4% stacking, 10% running gel). Then proteins were transferred to polyvinylidene fluoride membranes (Immobilon®-P Millipore), according to standard protocols. After blocking with 5% non-fat milk powder in 0,1% Tween 20 in Tris-buffered saline solution for 1h at room temperature (RT), membranes were incubated overnight at 4 °C with the primary antibodies described in the table below (Table 1.) diluted in blocking buffer. Blots were incubated for 2h at RT with the corresponding alkaline phosphatase-linked anti-mouse, anti-rabbit, or anti-goat secondary antibodies, as described in Table 1. The protein bands were visualized with enhanced chemiluminescence substrate (ECF, GE Healthcare) using chemifluorescence imaging (ChemiDocTM MP imaging system, Bio-Rad).

**Table 1.**
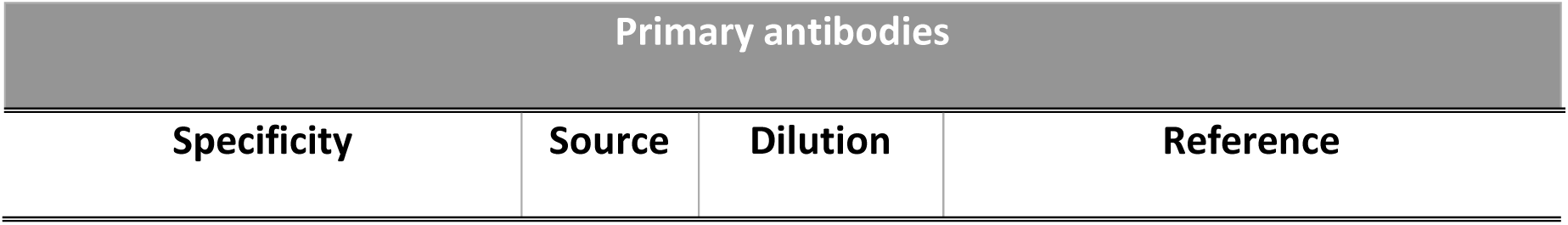
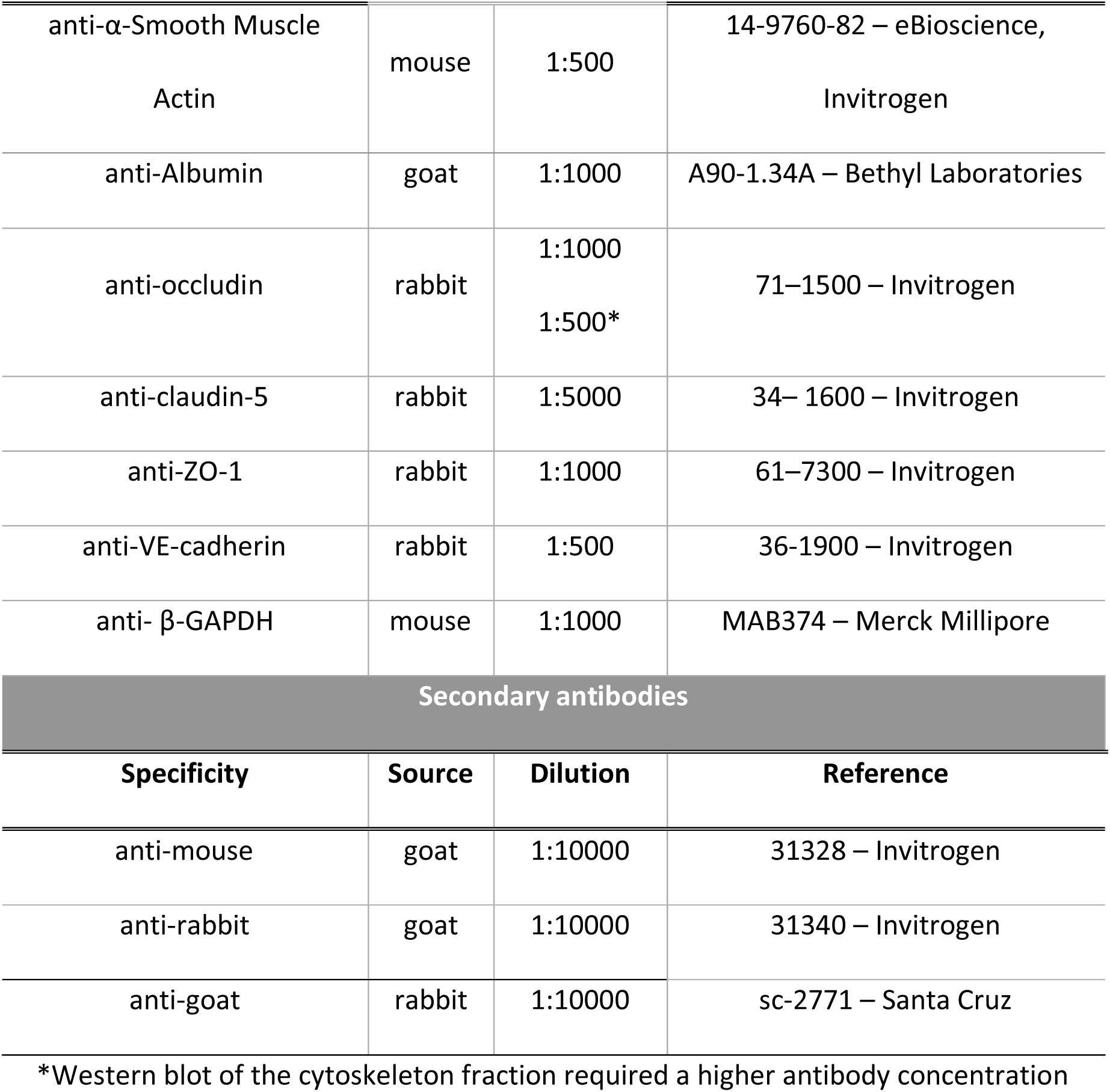
List of antibodies used for western blot analysis.

| Primary antibodies |  |  |  |
| --- | --- | --- | --- |
| Specificity | Source | Dilution | Reference |
| MSC for Neurovascular Repair in MJD |  |  |  |
| anti-α-Smooth Muscle<br>Actin | mouse | 1:500 | 14-9760-82 – eBioscience,<br>Invitrogen |
| anti-Albumin | goat | 1:1000 | A90-1.34A – Bethyl Laboratories |
| anti-occludin | rabbit | 1:1000<br>1:500* | 71–1500 – Invitrogen |
| anti-claudin-5 | rabbit | 1:5000 | 34– 1600 – Invitrogen |
| anti-ZO-1 | rabbit | 1:1000 | 61–7300 – Invitrogen |
| anti-VE-cadherin | rabbit | 1:500 | 36-1900 – Invitrogen |
| anti- β-GAPDH | mouse | 1:1000 | MAB374 – Merck Millipore |
| Secondary antibodies |  |  |  |
| **Specificity** | **Source** | **Dilution** | **Reference** |
| anti-mouse | goat | 1:10000 | 31328 – Invitrogen |
| anti-rabbit | goat | 1:10000 | 31340 – Invitrogen |
| anti-goat | rabbit | 1:10000 | sc-2771 – Santa Cruz |

\*Western blot of the cytoskeleton fraction required a higher antibody concentration

Semi-quantitative analysis based on the optical density of scanned membranes was carried out using ImageJ software (Fiji, ImageJ; version 1.51). The specific optical density was normalized to the amount of glyceraldehyde 3-phosphate dehydrogenase (GAPDH) or total protein loaded in the corresponding lane of each gel.

### Immunofluorescence

For standard immunohistochemical, sagittal sections of the left hemisphere of the cerebellum were cut with a thickness of 30 μm, using a cryostat (CryoStar NX50, Thermo Scientific, Thermo Fisher Scientific) at −19 °C. Sections were collected in anatomical series and placed directly onto super-frost microscope slides (Superfrost® plus, Thermoscientific). Slides were stored at −20 °C until further processing.

The immunohistochemical procedure was performed in sections previously collected and placed on superfrost slides. Firstly, sections were hydrated with PBS 1x for 30 min, followed by 1 hour of blocking and permeabilization at room temperature in 3% bovine serum albumin (BSA) with 0.1% Triton X-100. Sections were incubated overnight at 4 °C with primary antibodies diluted in 0.3% BSA: goat anti-Colllagen type IV (CoIV; 1:250, #AB769 Millipore), rabbit anti-Fibrin/FITC (1:40, #F011102-2 Dako) and rabbit anti-von Willebrand factor (vWF; 1:100, PA5-16634 Invitrogen). Sections were then washed with PSB and incubated for 2 hours at room temperature with the respective secondary antibody coupled to the fluorophores: anti-goat Alexa Fluor 568 (1:200, A11057 Thermo Fisher Scientific) or anti-rabbit 488 (1:200, A21206 Thermo Fisher Scientific), respectively. Nuclear staining was performed by incubation with DAPI (4′,6-diamidino-2-phenylindole, 2 µg/mL in PBS, Invitrogen) for 10 min. Finally, sections were washed and cover-slipped with Dako fluorescence mounting medium (S3023, Dako).

### Determination of blood vessel coverage: quantification of CoIV- and vWF-positive area/intensity

The CoIV-positive area was quantified in tile images of the cerebellum acquired with Plan-Apochromat 20X/0.8 M27 objective in Zeiss Axio Imager Z2 microscope with red (561 nm) laser. Three to four sagittal sections per animal (inter-section distance: 180 μm) corresponding to a similar cerebellar region were analyzed using the Image J software (National Institutes of Health; USA). The same fluorescence intensity threshold was applied to all images, and the percentage of surface area positive for CoIV staining in the cerebellum was measured.

Similarly, the vWF-positive area and intensity was quantified in tile images of the cerebellum acquired with a Plan-Apochromat 20X/0.8 M27 objective in ZEISS Axio Scan.Z1 Slide Scanner microscope with green (488 nm) laser. Three to four sagittal sections per animal (inter-section distance: 180 μm) corresponding to a similar cerebellar region were analyzed using QuPath software (v0.4.3) (Bankhead et al., 2017). The same fluorescence intensity threshold was applied to all images and the percentage of surface area positive for vWF staining in the cerebellum was measured.

### Evans Blue Dye Assay

#### Evans Blue injection and tissue collection

Mice were intravenously injected with a 2% solution of EB dye (100 µL/30 g body weight) via the caudal vein. After 30 min, the animals were deeply anesthetized intraperitoneally and sacrificed by transcardial perfusion with ice-cold PBS (pH 7.4). Perfused brains were harvested and cut in halves. Right hemispheres were use to quantify EB via spectrophotometric analysis. Left hemispheres of the brain were preserved in tissue-tek (Sakura) at −80 °C to be cut in slices and used for fluorescence microscopy. For more details, please see the schematic representation procedure in Figure 4.3A.

#### EB detection by spectrophotometry

For EB spectrophotometric analysis, the liver and the right hemisphere of the cerebrum and cerebellum were incubated overnight with pure formamide, using six times the volume of each organ’s weight. The incubation was performed under continuous agitation in a water bath at a temperature of 70°C. Afterwards, samples were centrifuged at maximum speed (4 °C, 20 min) and the supernatant’s absorbance was measured at 620 nm and 720 nm using the Fluorimeter SpectraMax Gemini EM (Molecular Devices). The final EB concentrations were calculated by subtracting absorbance values at 620 nm to absorbance values at 720 nm and compared with a pre-defined standard curve. Cerebellum and cerebrum values were normalized with liver EB concentration to exclude EB injections variability.

#### EB detection by fluorescence microscopy

Sagittal sections of the left hemispheres of the cerebella were cut with a thickness of 30 μm, and placed directly onto superfrost microscope slides (Superfrost® plus, Thermo Scientific). Slides were stored at −20 °C until further processing.

Fluorescence microscopy analysis was performed on brain sections (6 to 11 per animal) from lateral to central coordinates. After sections were hydrated with PBS 1x for 30 min, nuclear staining was performed by incubation with DAPI, and slides were cover-slipped on Dako fluorescence mounting medium. Cerebellar tile images were acquired with Plan-Apochromat 20X/0.8 M27 objective in ZEISS Axio Scan.Z1 Slide Scanner microscope with red (561 nm) and Blue (405 nm) filters. The number of EB deposition signals detected on each analyzed slide was counted as an individual event. The total number of events observed per animal was then determined.

### Behavioral Assessment

Animals of the 8-month-old set were subjected to motor tests before the first MSC injection and one week after the last treatment (according to the timeline represented in Figure 4.1 A). All tests were performed after acclimatization during the active period of the animals in a dark room lit with only red light. Animal cycles were inverted 48 h before behavioral testing. Beam-walking and footprint tests was used to assess motor behavior, coordination, and gait. A detailed description of the tests is provided in the Supplemental methods and materials.

### Statistical Analysis

All statistical analyses were performed using GraphPad Prism (version 9.0, GraphPad Software, La Jolla, CA, USA). Statistical differences were assessed by an unpaired Student’s t-test to compare means between two groups and one-way Analysis of Variance (ANOVA), followed by the adequate post-hoc Tukey test for multiple comparisons. In the ICG clearance assay and behavioral tests, the effect of group and/or time on all variables was assessed using repeated two-way ANOVA, followed by adequate post-hoc Tukey test to identify significant differences among groups and time periods. Pearson or spearman correlation coefficient was determined to study the relationship between two parametric or non-parametric variables, respectively. Outliers were removed according to Grubb’s test (alpha = 0.05). Results are expressed as mean ± standard error deviation (SD). P-values were considered statistically significant according to the following criteria: p > 0.05 = not significant; * p ≤ 0.05, ** p < 0.01, *** p < 0.001, and **** p < 0.0001.

## Results

### Repeated Systemic MSC administration alleviates MJD phenotype in Tg-ATXN3-69Q Mice, even in advanced disease stages

It is well established that Tg-ATXN3-69Q mice used in this study exhibit marked cerebellar atrophy, severe motor coordination deficits, and gait ataxia, which resemble the human MJD phenotype (8).

We had previously shown that four repeated MSC treatments improved motor deficits in young (1-2-month-old) MJD mice (15). Nonetheless, the effectiveness of the therapeutic approach adopted in the present study – two consecutive intravenous MSC treatments (spaced 1-2 weeks apart) to older MJD animals – in alleviating MJD motor impairments was not unequivocally certain. Thus, as a proof-of-concept of the therapeutic efficacy, motor performance was assessed by beam-walking, and footprint, one week before (week 0) and one week after (week 3) MSC treatment as illustrated in the timeline (Figure 1A).

First, gait deficits were assessed by analyzing footprint patterns. Regarding the left foot overlap measurements, Tg Nt mice had a higher left overlap distance when compared to controls (Figure 1B; week 3, Wt: 0.744±0.224 vs Tg Nt: 1.248±0.440, p=0.0004). Additionally, Tg T mice exhibited significantly decreased left foot overlap distance at week 3 compared to Tg Nt (Figure 1B; week 3 Tg Nt: 1.248±0.440 vs Tg T: 0.851±0.351 p=0.0245), with no differences observed between Wt and Tg T after treatment (Figure 1B; week 3 Wt: 0.744±0.224 vs Tg T: 0.851±0.351p= 0.7186).

Motor coordination and balance were then evaluated using the beam-walking test. Tg Nt mice took significantly longer to cross the beam than their Wt littermates due to motor deficits (Figure 1C; week 3 Wt: 2.268±0.512 vs Tg Nt: 16.566±3.184, p<0.0001). Additionally, Tg T mice showed a weak tendency to improve compared to Tg Nt animals (Figure 1C; week 3 Tg Nt: 16.566±3.184 vs Tg T: 14.786±2.372; p=0.1055).

Given the potential for vascular abnormalities in the Tg-ATXN3-69Q mice (9), we performed a comparative evaluation of MSC biodistribution in MJD and Wt animals. For that, 8-month-old mice received an IV injection of MSC expressing luciferase (MSC-Luc) followed by an IP injection of luciferin. Luciferase is an enzyme that degrades luciferin, producing a bioluminescent signal. The bioluminescent signal was captured 30 min after MSC administration, focusing on the cranial region (Sup Figure 1A, B). MSC-luc reached the brains of both Wt and Tg animals, and although Tg had a lower signal detected compared to Wt animals, this decrease was not statistically significant (Sup Figure 1C; Wt T: 3.102±1.428 vs Tg T: 2.206±1.209; p=0.1358).

Taken together, these results suggest that, surprisingly, two MSC treatments can alleviate gait deficits in more advanced stages of MJD. Moreover, the bioluminescent assay suggests that the poor biodistribution induced by vascular abnormalities may not be a significant factor hindering the therapeutic effects of MSC in the brain. Upon confirming the therapeutic strategy’s effectiveness in ameliorating MJD phenotype, we then evaluated cerebrovascular impairments and MSC treatment on them.

### Blood Vessel function is impaired in Tg-ATXN3-69Q mice

Impairments in cerebrovascular structure and function emerged as significant clinical hallmarks of neurodegenerative diseases, including PolyQ disorders (10, 12, 25, 26). These findings highlight the important role of NVU dysfunction in the progression of neurodegenerative disorders. To investigate whether similar alterations are present in MJD, we evaluated potential functional impairments in blood vessels in Tg-ATXN3-69Q mice. To assess changes in cerebral blood volume and clearance, we performed an experiment in which 7-month-old animals received an IV injection of the ICG contrast agent via the tail vein. Upon injection, ICG can be present in the bloodstream as a free molecule (775 Da) or can bind to proteins like albumin, forming higher molecular weight complexes. Real-time ICG fluorescent signal in the cerebellum was then measured 10, 30, and 60 min after injection (procedure summarized in Figure 2A).

**Figure 2.**
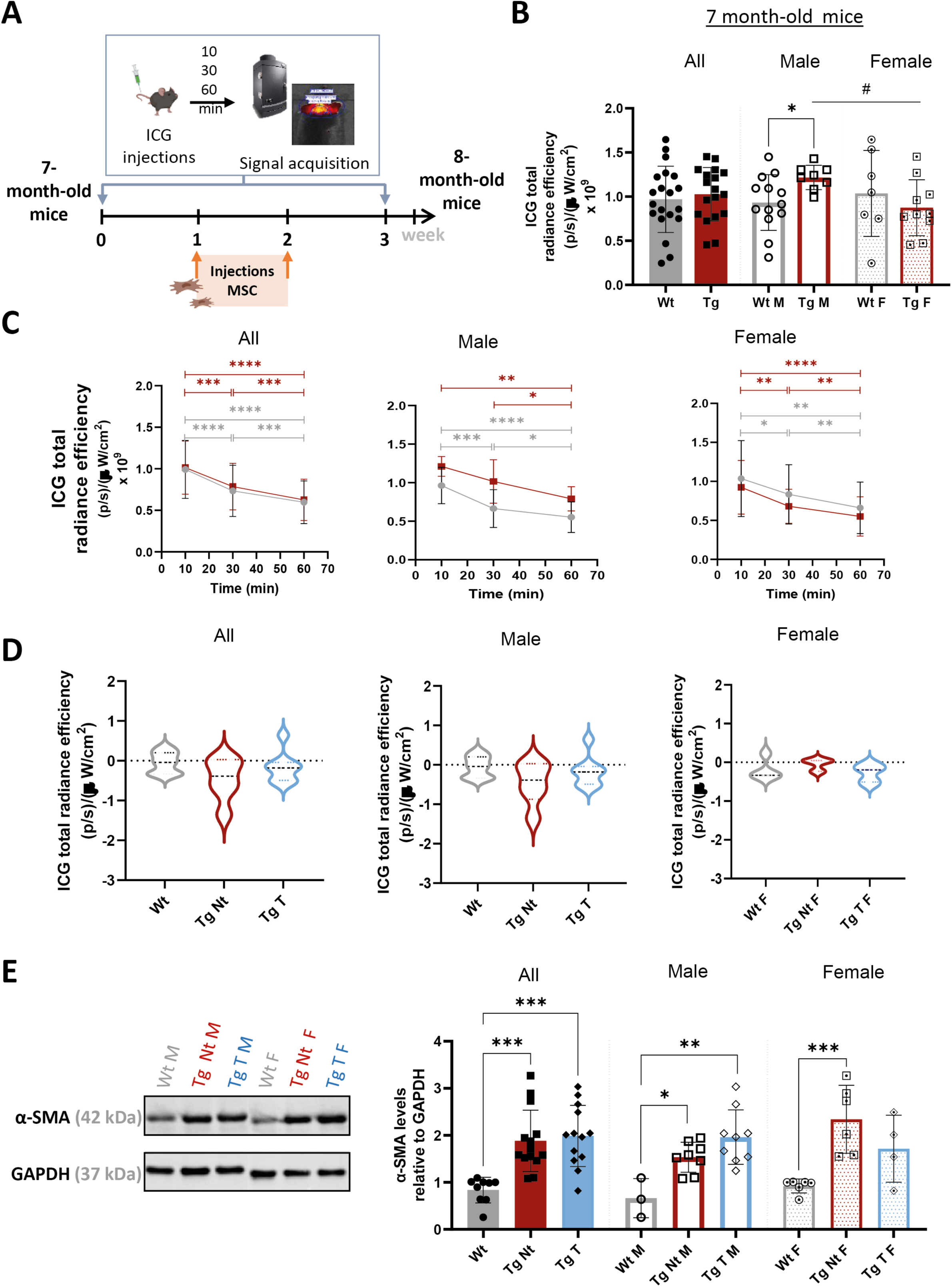
Evidence of blood vessel functional impairments in Tg-ATXN3-69Q mice and MSC treatment partially revert them. Schematic representation of the Indocyanine Green (ICG) experiment (A): mice received an IV injection of the contrast agent ICG, and signal intensity of ICG was measured in the cerebellum, using one week before (week 0) and one week after (week 3) treatment an IVIS equipment. ICG signal intensity was detected in the cerebellar region of mice (Wt n=20 [13♂+7♀] vs Tg n=18 [8♂+10♀]) 10 min upon injection ICG at week 0 (B). Clearance of ICG after injection (C): cerebellar ICG signal intensity assessed 10, 30, and 60 min in mice (Wt n=8 [4♂+4♀] vs Tg Nt n=6 [3♂+3♀] vs Tg n=8 [3♂+5♀]) upon injection at week 0. The variation of the ICG signal measured in the cerebellum of animals (Wt n=20 [13♂+7♀] vs Tg n=18 [8♂+10♀]) before (week 0) and after (week 3) MSC treatment was evaluated (D) through the calculation of the longitudinal ICG differences: ICG signal week 3 - ICG signal week 0. Representative WB membrane and quantification of the expression levels of α-SMA in the cerebellum 8-month-old mice (Wt n=10 [3♂+7♀] vs Tg Nt n=14 [8♂+6♀] vs Tg T n=13 [9♂+4♀]) (E). Protein relative levels were normalized with GAPDH. Data presented as mean ± SD. Unpaired t-test, two-way ANOVA followed by Tukey’s test or one-way ANOVA followed by Tukey’s test. p>0.05 = not significant; * p<0.05, ** p<0.01, ***p< 0.001, ****p< 0.0001.

Our results revealed an increase in ICG signal in the cerebellum of 7-month-old male Tg animals compared to Wt, 10 min post-injection (Figure 2B; Wt M: 0.8317×109±0.3847 vs Tg M: 0.5332×109±0.3249), while no alterations were observed in female Tg mice when compared to Wt (Figure 2B; Wt F: 1.036×109±0.1372 vs Tg F: 0.8740×109±0.3164; p=0.4165; Tg M vs Tg F p=0.0118). Additionally, we evaluated ICG clearance by comparing fluorescent signals at 10-, 30-, and 60-min post-injection in both Wt and Tg mice. The 7-month-old male Wt and Tg mice displayed different ICG clearance profiles. Wt male mice exhibited an ICG signal decrease of 31% from the 10 to the 30-minute time point (Figure 2C; Wt M 10 min: 0.962×109±0.235 vs Wt M 30 min: 0.663×109±0.245; p=0.001) and a smaller decrease of 11.4% from the 30 to the 60 min time point (Figure 2C; Wt M 30 min: 0.663×109±0.245 vs Wt M 60 min: 0.553×109±0.198; p=0.0424), resulting in an overall decrease of 42.6% reduction from 10 to 60 min post-ICG injection. In contrast, Tg male animals demonstrated a slower clearance rate of ICG signal, with no significant alterations observed from the 10 to the 30-minute time point (Figure 2C; Tg M 10 min: 1.211×109±0.128 vs Tg M 30 min: 1.015×109±0.282; p= 0.2173), followed by an 18,6% ICG signal reduction between the 30 to the 60-minute time point (Figure 2C; Tg M 30 min: 1.015×109±0.282vs Tg M 60 min: 0.7904×109±0.15; p= 0.0434), leading to a decrease of 34.7% reduction from 10 to 60 min post-ICG injection. These results suggest that there is a potential impairment in blood clearance in Tg mice compared to Wt animals, though these alterations may be influenced by sex.

Interestingly, dysregulation in cerebellar blood volume and clearance was also detected in 2.5-month-old mice (Sup. Figure 2 A, B), although at that age, blood volume was decreased in Tg mice compared to controls. These results suggest that such dysregulation is disease-stage dependent.

To investigate whether MSC treatment could alleviate these dysregulations, animals received two consecutive intravenous MSC treatments (spaced 1-2 weeks apart) and the fluorescent ICG signal was detected in mice cerebella 30 min ICG post-injection. the variation in the ICG signal measured before and after treatment was determined (longitudinal ICG differences = ICG signal week 3 - ICG signal week 0).

In males, Wt mice only demonstrated minimal changes in ICG signal, while Tg Nt mice displayed a decrease during the experiment, presenting a significantly different profile compared to controls (Figure 2D; Wt M: 0.0949±0.1826 *vs* Tg Nt M: −0.8740±0.4260, p=0.0394). MSC administration appeared to prevent ICG signal variation, as we noted a tendency to diminish the ICG variation degree/extension in Tg T mice compared to Tg Nt animals (Figure 2D; Tg Nt M: - 0.8740±0.4260 *vs* Tg T M: −0.0054±0.5891, p=0.0784) and no significant differences between Wt and Tg T profiles were observed (Figure 2D; Wt M *vs* Tg T M p=0.9446). No significant differences were detected between female groups.

To assess blood vessel contractility capacity, the expression levels of α-smooth Muscle Actin (α-SMA) were determined by western blotting (as an indirect evaluation). α-SMA, a cytoskeletal protein that is highly expressed in brain vessels, plays an important role in vessel constriction (27). Notably, there was a tendency significant increase in α-SMA levels in Tg Nt compared to Wt (Figure 2E; Wt: 0.1926±1.094 *vs* Tg Nt: 1.080±3.267, p=0.0002). A tendency to this increase was already noticeable in 2.5-month-old Tg mice (Sup Figuer 1C). The administration of MSC may have had opposite effects on males and females. In males, Tg T mice exhibited a greater increase in α-SMA expression than their Tg Nt counterparts when compared to Wt (Figure 2E; Wt M: 0.6651±0.4183; Tg Nt M: 1.536±0.3198; Tg T M: 1.963±0.5784; Wt M *vs* Tg Nt M: p= 0.0353; Tg Nt M *vs* Tg T M: p=0.0018). Conversely, in females, Tg Nt mice exhibited a significant increase in α-SMA expression compared to Wt (Figure 2E; Wt F: 0.8188±0.3073 *vs* Tg Nt F: 2.342±0.7227, p= 0.0009), while Tg T display only a tendency to increase α-SMA levels relative to control animals (Figure 1; Tg Nt F: 2.342±0.7227 *vs* Tg T F: 1.717±0.7106; Tg Nt M *vs* Tg T M: p=0.0650).

Altogether, these results suggest that MSC therapy can ameliorate some of the functional vascular abnormalities observed in 8-month-old MJD mice of the Tg-ATXN3-69Q model and may have a sex-specific effect.

### MSC administration reverts vascular coverage alterations in Tg-ATXN3-69Q mice

Given the functional impairments in cerebellar vasculature detected, we also investigated potential structural vascular alterations. For that we quantified blood vessel coverage, within the cerebellum by analyzing the surface area positive for CoIV, a marker of blood vessels’ basement membrane. Tg mice showed a tendency to increase CoIV surface area compared to controls (Figure 3A-B; Wt: 10.14±1.495 vs Tg: 13.82±3.305; p=0.0877). Notably, when we analyzed data by sex, it became evident that Tg females (Tg F) exhibited no alterations, while a significant increase in CoIV surface area was observed in 8-month-old Tg males (Tg M) compared to Wt littermates. MSC treatment significantly decreased the CoIV surface area in the cerebella of Tg T mice in comparison to Tg Nt animals (Figure 3B; Tg Nt M: 16.50±0.6701 *vs* Tg T: 10.04±0.2486, p<0.0001).

**Figure 3.**
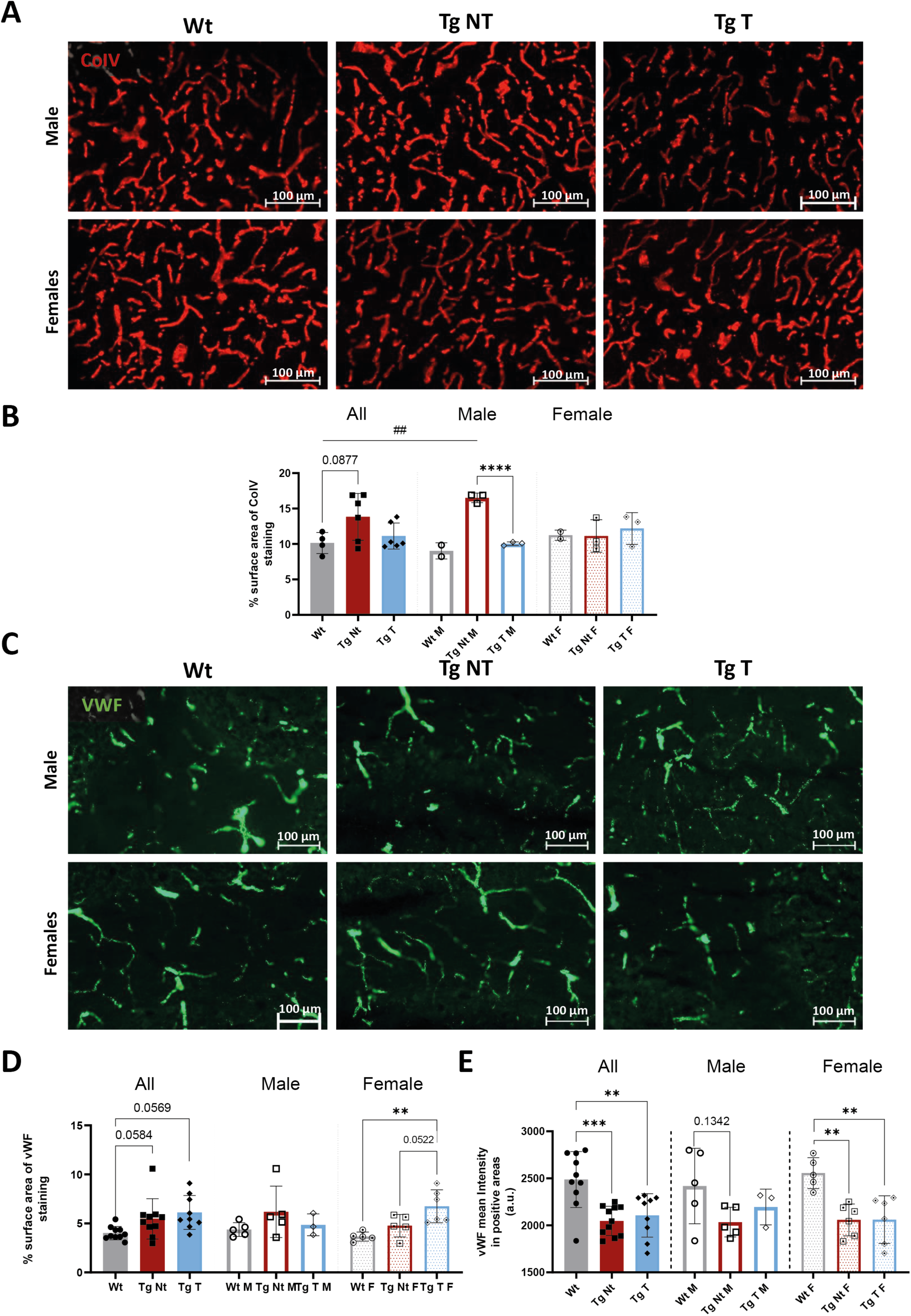
MSC partially ameliorated cerebellar dysregulation in vascular coverage in 8-month-old Tg-ATXN3-69Q mice. Representative images (A) and quantification (B) of the surface area of CoIV staining in the cerebella of Wt, non-treated (Tg Nt) and MSC-treated Tg mice (Wt n=4 [2♂+2♀] vs Tg Nt n=6 [3♂+3♀] vs Tg T n=6 [3♂+3♀]). Representative images (C) and quantification of the surface area (D) and intensity (E) of vWF staining in the cerebella of mice (Wt n=10 [5♂+5♀] vs Tg Nt n=10 [5♂+5♀] vs Tg T n=9 [3♂+6♀]). Data presented as mean ± SD. Unpaired t-test test or one-way ANOVA followed by Tukey’s test. p>0.05 = not significant; ** p<0.01, ***p< 0.001, ****p< 0.0001.

Additionally, we evaluated alterations in vascular endothelial cell coverage by assessing surface area positive for vWF, an endothelial cell marker. We observed a tendency for an increase in vWF coverage in 8-month-old Tg mice in both males and females (Figure 3D), which was significant in 2.5-month-old Tg mice (Sup. Figure 2E). Furthermore, we evaluated the levels of vWF in the vascular endothelial cells to determine the presence of vascular injury. vWF is a multimeric glycoprotein that in the brain is produced by vascular endothelial cells and stored in these stimuli cells in Weibel-Palade bodies (28). In response to an insult like increased TNF-α levels, endothelial cells release the contained Weibel-Palade bodies towards the lumen of blood vessels and simultaneously decrease mRNA and protein expression of vWF (29). Consequently, this results in reduced vWF levels within endothelial cells. Here we observed a decrease in the mean intensity of vWF in cerebellar vascular endothelial cells of 8-month-old Tg mice compared to controls (Figure 3E; Wt: 2487± 298 vs Tg: 2047±156.2; p=0.0004). Treatment had no effect in altering vWF coverage or intensity.

Altogether, these results suggest that there is an increase in cerebellar blood vessel coverage in MJD Tg mice and the presence of vascular injury. MSC administration could only partially revert this alteration.

### MSC restores BBB leakage in the cerebella of 8-month-old Tg-ATXN3-69Q mice

BBB breakdown is a key feature of MJD. Indeed, as previously demonstrated by Lobo et al., BBB permeability to large molecules is increased in 16/17-month-old Tg-ATXN3-69Q mice (9). Interestingly, here we found evidence of this increased permeability in 8-month-old Tg mice but not in younger 2.5-month-old animals (Sup. Figure 3).

To evaluate whether MSC treatment could ameliorate BBB leakage, we performed an EB dye assay, a widely used method to evaluate BBB disruption. Upon IV injection, EB strongly binds to blood albumin, forming a complex of approximately 69 kDa that cannot cross the BBB under physiological conditions. However, when BBB integrity is compromised, the EB-albumin complex accumulates in the brain parenchyma (30). Thus, in our experiment, 8-month-old Wt, non-treated, and MSC-treated Tg male mice received an IV injection of EB in the caudal vein. Thirty minutes later, mice were intracardially perfused with PBS to remove EB from circulation, and brains were harvested (timeline is represented in Figure 4A).

**Figure 4.**
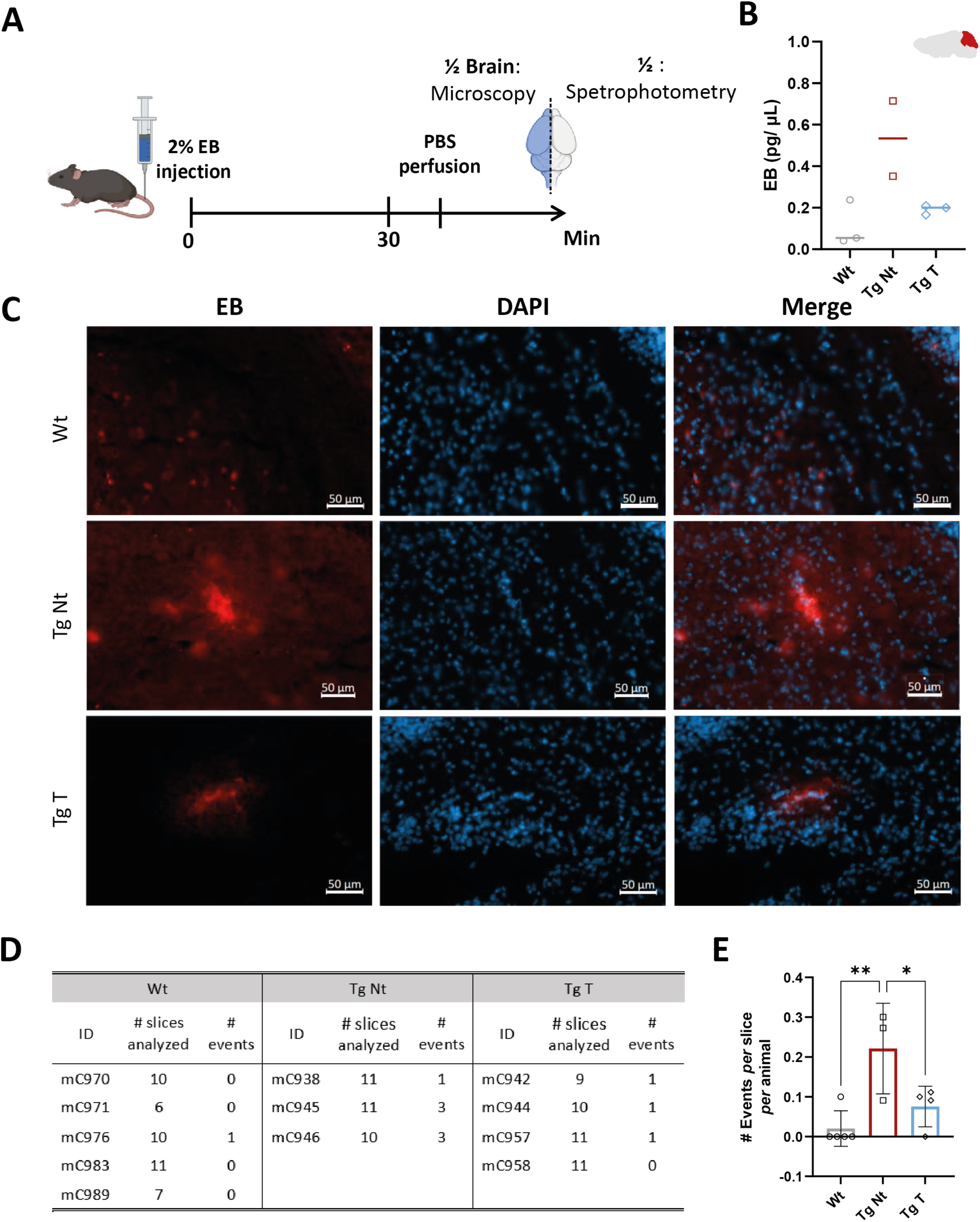
MSC can partially restore BBB leakage in 8-month-old Tg-ATXN3-69Q mice. Schematic representation of the Evans Blue dye (EB) experiment (A): 8-month-old mice were injected with 2% EB in the tail vein. EB concentration (pg/μL) determined by spectrophotometry (Wt n=3 vs. Tg Nt n=2 vs. Tg T n=3) (B). Representative fluorescence images of EB extravasation in the cerebellum of Wt, non-treated (Tg Nt), and MSC-treated (Tg T) Tg male mice (C). Table summarizing the number of individual events (EB deposition signals) detected and the number of analyzed sections per animal (D). Graphical representation of the total number of events observed per analyzed section of each animal (Wt n=5 vs Tg Nt n=3 vs Tg T n=4) (E). Representative WB membrane and quantification of Albumin protein levels in the cerebellum of mice (Wt n=13 [4♂+9♀] vs Tg Nt n=14 [6♂+8♀] vs Tg T n=15 [6♂+9♀]) (F). *Data presented as mean ± SD. One-way ANOVA followed by Tukey’s test. p>0.05 = not significant; * p<0.05, ** p<0.01*.

The concentration of EB in the cerebellum was determined via spectrophotometry, and values were normalized to liver EB concentrations to exclude variations in the IV procedure. Non-treated Tg animals exhibited higher EB levels compared to Wt littermates (Figure 4B; Wt: 0.1110±0.1101 *vs* Tg Nt: 0.5336±0.2563). Additionally, Tg T mice display lower EB concentrations when compared Tg Nt mice (Figure 4B; Tg Nt: 0.5336±0.2563 *vs* 0.1920±0.02242). No statistical analyses were performed due to a low number of Tg Nt subjects.

Moreover, the presence of EB extravasation was confirmed using fluorescence microscopy (Figure 4C). Brain sections spanning from lateral to central coordinates were evaluated and each EB deposition signal detected in the cerebellum *per* slice was counted as an individual event of BBB leakage. The cumulative number of events per animal was then calculated. Details of this assessment are described in Figure 3D. As expected, the number of EB extravasation events *per* slice *per* animal was higher in the cerebellum of Tg Nt mice than in Wt littermates (Figure 4E; Wt: 0.0200±0.0447 *vs* Tg Nt: 0.2212±0.1137, p=0.0073). Importantly, there were fewer EB depositions detected in Tg T mice than in Tg Nt animals (Figure 4E; Tg Nt: 0.2212±0.1137 *vs* Tg T: 0.0755±0.0510, p=0.0490), suggesting that MSC administration could partially restore BBB integrity.

### Administration of MSC ameliorates AJ/TJ-associated protein delocalization

Considering the critical role that AJ/TJ-associated proteins play in the regulation of BBB permeability, we evaluated a potential dysregulation in the expression levels and subcellular location of such proteins. Moreover, we investigated whether MSC could modulate AJ/TJ protein levels in the subcellular location where they fulfill their barrier function.

Our results showed a tendency towards a decrease in total expression levels of the transmembrane protein occludin in 0.5-month-old tg mice compared to wt animals, which became significant with MJD progression (Sup. Figure 4 A-C). Regarding occludin subcellular location, in 8-month-old Tg mice, there was a reduction in levels of membrane fraction compared to controls (Figure 5B; 1± 0.1812 vs. Tg: 0.5691±0.1517; p<0.0001), with no alterations observed in the cytoplasmic fraction (Figure 5A). Notably, MSC treatment partially restored occludin levels in the membrane subfraction, but only in females (Figure 5B; Tg Nt F: 0.5630±0.1438 vs Tg T F: 0.7866±0.1525, p=0.0330).

**Figure 5.**
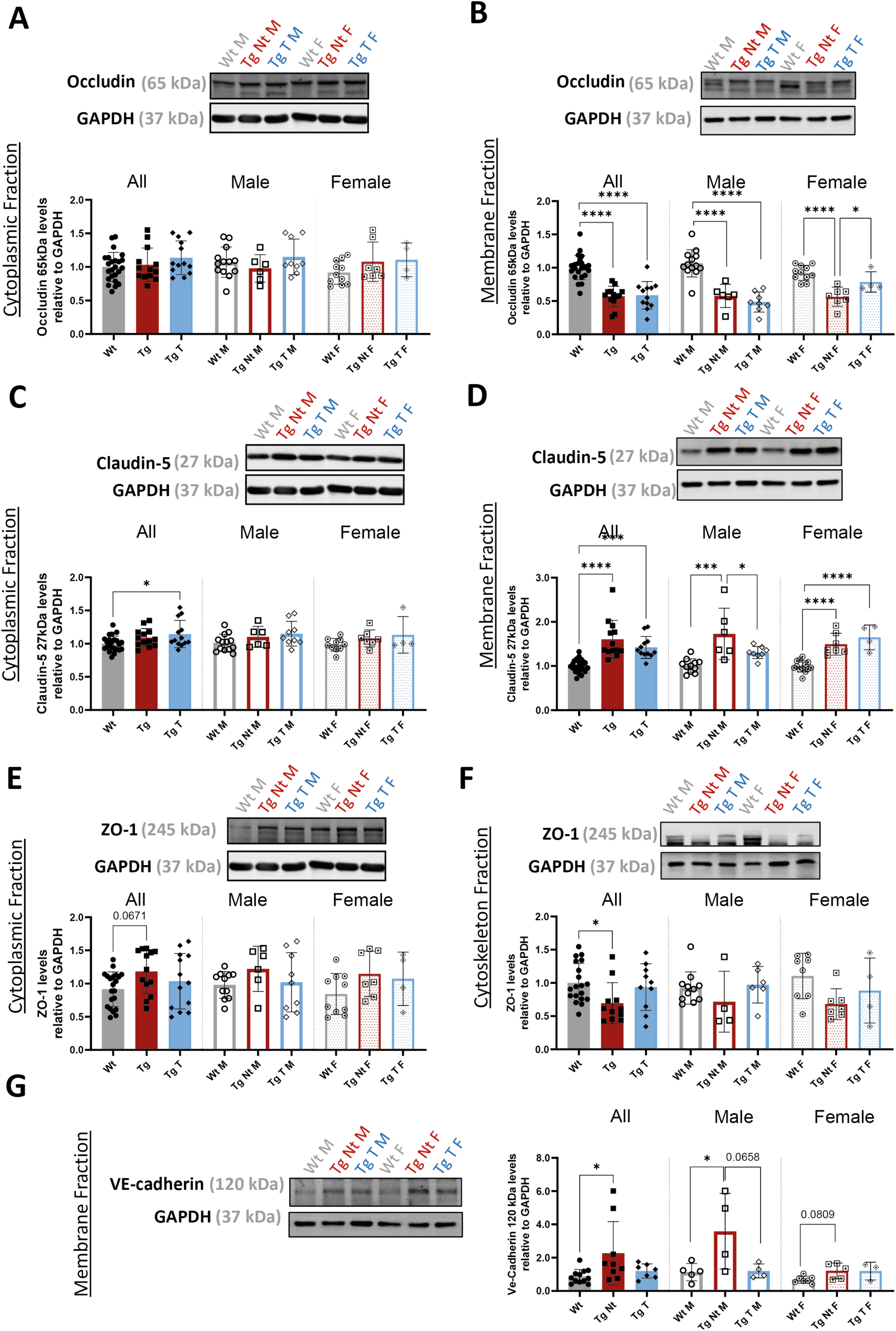
MSC treatment partially restores AJ/TJ levels, in a sex-specific way, in Tg-ATXN3-69Q Mice. Representative WB membranes and quantification of the expression levels of Occludin 65 kDa isoform in cytoplasmatic (A) and membrane (B) protein extracts obtained from the cerebella of 8-month-old mice, Claudin-5 27 kDa isoform in cytoplasmatic (C) and membrane (D) protein extracts (Wt n=24 [13♂+7♀] vs Tg Nt n=13 [6♂+7♀] vs Tg T n=12 [8♂+4]), ZO-1 in cytoplasmatic (E) and cytoskeleton (F) protein extracts (Wt n=19 [10♂+9♀] vs Tg Nt n=11 [4♂+7♀] vs Tg T n=10 [6♂+4♀]) and VE-Cadherin in membrane protein extracts (G) (Wt n=12 [5♂+7♀] vs Tg Nt n=9 [4♂+5♀] vs Tg T n=7 [4♂+3♀]). Data presented as mean ± SD. One-way ANOVA followed by Tukey’s test. p>0.05 = not significant; * p<0.05, ***p< 0.001, ****p< 0.0001.

In contrast, there was an increase in the levels of the claudin-5 27 kDa monomer in the cerebellum of 2.5- and 8-month-old Tg mice when compared to Wt (Sup. Figure 4 D-J). This increment was consistently observed in the cytoplasmic and membrane protein extracts of 8-month-old mice (Figure 5C; cytoplasm - Wt: 1±0.1142 vs Tg: 1.090±0.1367; p=0.0414; Figure 5D; membrane - Wt: 1±0.1384 vs Tg: 1.507±0.2820; p<0.0001). MSC administration significantly decreased claudin-5 levels in Tg T male animals relative to non-treated Tg littermates (Figure 5D; Tg Nt M: 1.726±0.5835 vs Tg T M: 1.306±0.1367, p=0.0387).

Furthermore, the total expression levels of the cytoplasmic adaptor protein ZO-1 decreased, but only in 8-month-old Tg mice, relative to controls (sup. Figure 4 K-M). Intriguingly, there was a tendency towards an increase in ZO-1 levels at the cytoplasmic level in 8-month-old Tg Nt mice relatively to Wt mice (Figure 5E; Wt: 0.9148±0.2614 vs Tg: 1.181±0.3294; p=0.0671), while a reduction was observed in Tg Nt animals in the cytoskeletal fraction (Figure 5F; Wt: 1±0.294 vs Tg: 0.6948±0.3087; p=0.0370). These suggest that not only ZO-1 expression but also its location and association with the cytoskeleton are impaired in MJD. MSC administration partially restored ZO-1 levels, as no differences were detected between Tg T and Wt mice (Figure 5E; Wt: 1±0.294 vs Tg T: 0.9374±0.3523, p=0.8659).

In addition to TJ proteins, adherent junction proteins, such as the transmembrane glycoprotein VE-cadherin also contribute to endothelial integrity and vascular permeability modulation (7, 31). Our results show a significant increase in 120 kDa VE-cadherin form total expression levels in the cerebellum of 8-month-old Tg mice compared to their Wt littermates (Sup. Figure 5). This increase was also observed in the membrane protein extracts (Figure 5G; Wt: 0.8671±0.4205 vs Tg: 1.343±0.5386; p=0.0461). Interestingly, the 120 kDa isoform was not detected in the cytoplasm fraction, suggesting that glycosylation primarily occurs in the cellular membrane. Moreover, MSC-treated Tg male mice showed a tendency to decrease the membrane levels of this AJ protein compared to Tg Nt animals (Figure 5G; Tg Nt M: 3.586±2.275 vs Tg T M: 1.203±0.4233, p=0.0658). In fact, no significant differences were detected in VE-cadherin expression between Tg T and Wt male mice (Figure 5G; Wt M: 1.132±0.5208 vs Tg T M: 1.203±0.4233, p=0.9964).

We also evaluated whether MSC could modulate AJ/TJ protein levels in total cerebellar protein extract, though no significant improvements were observed (data not shown).

In addition, we assessed the correlation between TJ subcellular localization levels and motor performance, as well as gait performance (Sup. Figure 5). Notably, we observed a strong correlation between motor performance in the beam walking test and ZO-1 levels in cytoskeleton protein extracts (Sup. Figure 5C; r=-0.7721, p=0,0033). Three different populations of animals could be distinguished - Wt, Tg Nt, and Tg T mice, with the Tg T population slightly converging towards the values of the Wt population.

Overall, our findings suggest that TJ/AJ protein levels and localization are compromised in MJD, albeit with disease-stage-dependent variations. Importantly, MSC could partially restore AJ/TJ protein re-localization, at least at the cellular membrane level. Thus, MSC protective action in the BBB possibly contributes to MJD pathology improvements

## Discussion

Although considerable research has been done on cerebrovascular and BBB dysfunctions in neurodegenerative disorders, the knowledge on these aspects in the context of MJD remains limited, with only one prior study addressing BBB impairment in MJD (9). In the present study, we demonstrate that cerebral vasculature is disrupted in a MJD mouse model (Tg-ATXN3-69Q (8)) marked by noticeable changes in blood vessel morphology and function, along with alterations in BBB permeability. Additionally, we provided compelling evidence of MSC’ ability to mitigate vascular abnormalities and BBB disruption in Tg-ATXN3-69Q mice. Specifically, our results revealed that the administration of MSC led to motor deficit ameliorations, which was concurrent with the alleviation of vessel density, and blood volume stabilization. Furthermore, our findings suggest that MSC modulation of AJ/TJ proteins contributed, at least in part, to a notable reduction in BBB disruption.

Over the last decade, accumulating evidence has underscored cerebrovascular dysregulation as a hallmark of neurodegenerative disorders including polyQ disorders (25). Within the context of MJD, it has been shown that patients have compromised cerebral blood flow (10–12) and higher BBB permeability, characterized by increased fibrinogen extravasation and concomitant microgliosis (9). Considering that these vascular alterations, namely BBB leakage, can exacerbate neuronal damage, the cerebrovascular unit has emerged as a potential therapeutic target, plausibly capable of modifying the progression of neurodegenerative disorders.

MSC are a promising cell therapy for the treatment of several neurodegenerative diseases and can modulate BBB breakdown (23, 32). Regarding PolyQ disorders, MSC have been shown to promote neuroprotection by enhancing cell survival, inducing neurogenesis, suppressing apoptosis, and regulating the immune system across diverse disease models (14). In line with preclinical studies, clinical trials have demonstrated that MSC are safe and may even delay the progression of these conditions. So far, 10 clinical trials, which are either completed or ongoing, aimed to test MSC safety and effectiveness in the treatment of PolyQ disorder (ClinicalTrials.gov: NCT02728115, NCT03252535, NCT04219241, NCT03378414, NCT01360164, NCT01489267, NCT01649687, NCT02540655, plus (18, 20)). Moreover, three preclinical studies have demonstrated that MSC-based therapies can promote phenotypical and neuropathological ameliorations in MJD Tg mouse models; however, sustained beneficial effects are only achieved with repeated administrations (15–17). Accordingly, in the present study, we found that two consecutive MSC injections ameliorated motor and gait impairments in 8-month-old Tg-ATXN3-69Q mice. The positive outcomes in 8-month-old MJD mice highlight the efficacy of MSC treatment even at later stages of the disease, given this model exhibits a severe ataxic phenotype.

Nevertheless, the exact mechanics through which MSC exert their protective action in MJD is still not fully understood, nor their potential effect on the vasculature. To the best of our knowledge, the present work is the first to show the potential of MSC to ameliorate neurovascular and BBB dysfunction in the context of a PolyQ disorder.

In the present study, we evaluated blood vessel functionality in Tg-ATXN3-69Q through an in vivo experiment where animals were injected with the ICG contrast agent. The variation in signal intensity of ICG represents its circulation in the bloodstream which was recorded in the cerebellum region. From the three time points assessing ICG signal, it reached its peak 10 min post-injections. At 7 months of age, Tg mice displayed a significant increase in ICG signal, suggesting that later cerebellar blood volume. Additionally, the curve variations of ICG signal intensity (from the 10- to 60-min time-point) evidenced the slower rate of blood clearance in Tg animals compared to Wt, which can be indicative of blood clearance impairments and/or ICG accumulation in brain parenchyma due to BBB leakage. Indeed, Lobo *et al*., show that older 16-17.5-month-old Tg-ATXN3-69Q also displayed greater cerebellar blood volume and BBB impairments relative to Wt (9). Accordingly, increases in cerebral blood volume have been observed in HD patients and mouse models, and such increment was accompanied by augmented vessel density (26, 33).

Given that α-SMA, a cytoskeletal protein primarily expressed in vascular smooth muscle cells and some pericytes (34), plays a key role in vascular contractility and the regulation of blood pressure and flow (27, 35), we assessed its expression levels in MJD mice. Our findings revealed an increase in α-SMA expression in the cerebellum of these animals, suggesting a potential contribution to the vascular dysfunctions observed in MJD. Similarly, a pre-clinical study demonstrated increases in α-SMA protein and mRNA levels in a mouse model of AD (36). Hutter-Schmid and Humpel proposed that such dysregulation could be a potential compensatory mechanism in response to impaired blood flow (36). Previous reports have highlighted reductions in cerebral blood flow in spinocerebellar ataxias, including in MJD patients, affecting mainly the cerebellum, but also the midbrain of the brainstem (10–12). Indeed, reductions in cerebral blood flow were found to be correlated with disease severity in SCAs, indicating that it may be a plausible disease biomarker (10, 12).

Moreover, Drouin-Ouellet *et al*. hypothesized that potential reductions in blood flow and oxygen delivery within the affected brain regions could lead to an increase in vascular density as a compensatory mechanism. Notably, the immunostaining for CoIV (basement membranes marker) and vWF (ECs marker) suggests an increase in cerebellar blood vessel density in the cerebellum of MJD 8-month-old Tg mice compared to Wt counterparts, which is in agreement with prior observations reported by our group in mice of this model with 16-17.5-months of age (9) and may in part explain the augmented cerebellar blood volume. A similar increase in vessel density has also been described in an HD mouse model as well as in HD patients (26, 37). Interestingly, treatment with MSC was able to modulate vascular coverage, as evidenced by the reduction in collagen IV surface area in MSC-treated Tg mice compared to untreated littermates, and the regulation of blood vessel function, influencing both blood volume and vessel contractility.

Furthermore, MSC administration led to a decrease in EB extravascular accumulation in the cerebellum of 8-month-old Tg male mice compared to non-treated animals, suggesting that MSC could alleviate BBB leakage. This diminishment in BBB permeability could be attributed, at least in part, to MSC restoration of TJ/AJ-associated proteins subcellular localization. Both TJ-associated proteins, such as occludin, claudin-5, and ZO-1, and AJ proteins, such as VE-cadherin, play vital roles in maintaining the stability of endothelial connections and regulating paracellular permeability in brain blood vessels. Thus, TJ/AJ are essential for BBB integrity (31, 38, 39). We found that demonstrated that AJ/TJ-associated proteins are dysregulated long before severe breakdown of the blood-brain barrier was detectable MJD Tg-ATXN3-69Q mice.

Here, we show that occluding expression levels were decreased in Tg-ATXN3-69Q mice in mice as young as 2 weeks of age. Although such dysregulation was not detected in older Tg-ATXN3-69Q 16-17.5-month-old mice (9), our results corroborate the literature reports regarding loss of occludin (65 kDa) in neurological disorders characterized by BBB breakdown, including in HD (26, 40). Additionally, upon evaluation of occludin distribution in subcellular compartments, we detected a decrease of occludin in the membrane fraction of 8-month-old MJD Tg-ATXN3-69Q mice. Being a TJ transmembrane protein, this membrane displacement indicates a loss of proper function by occludin in MJD mice. Contrarily, we found that monomeric claudin-5 form (27 kDa) expression levels were increased in the Tg-ATXN3-69Q model, even at 2.5 months of age, and in the cytoplasm and membrane fraction of 8-month-old MJD mice. Such results align with Lobo et al. previous report of increased monomer claudin-5 expression in 16-17.5-month-old Tg-ATXN3-69Q mice (9). Indeed, it has been proposed that an increment in claudin-5 expression may represent a compensatory mechanism in light of Occludin degradation (41, 42). We also detected the presence of a 100 kDa oligomeric claudin-5 form, believed to be the oligomerized claudin-5 within the cytoskeleton (43). This suggests that claudin-5-cytoskeleton assembly, crucial to the proper arrangement of TJ structure (43), is compromised in MJD. In parallel, we also evaluated the expression levels of the cytoplasmic adaptor ZO-1, responsible for occludin and claudin-5 binding to actin cytoskeleton (44). We show that ZO-1 total expression levels were decreased in the cerebellum of 8-month-old MJD Tg-ATXN3-69Q mice, similar to the fundings in other neurodegenerative disorders such as AD (45). However, we observed a reduction of ZO-1 levels in the cytoskeletal cerebellar fraction of Tg-ATXN3-69Q mice, despite its increase in its levels at the cytoplasmic level in 8-month-old Tg mice. These results indicate a possible delocalization of ZO-1 from the cytoskeleton and consequent accumulation in the cytosol in the cerebellum of these MJD mice, which could contribute to hamper TJ stability.

Apart from the TJ, AJ proteins also contribute to the formation and maintenance of BBB functions. Thus, we evaluated the expression levels and cellular location of VE-cadherin, an important AJ transmembrane glycoprotein of approximately 120 kDa. In the brain, VE-cadherin is specifically expressed by endothelial cells, and it is required to stabilize endothelial cell-cell interactions (39). VE-cadherin decreased expression has been reported in both acute neuronal disorders, like traumatic brain injury (46), and chronic neurodegenerative disorder, namely in AD patients’ brain vessels and mouse models (47)and in HD iPSC-Derived Brain Microvascular Endothelial Cells (48, 49). In contrast, we observed an increase in VE-cadherin expression levels in the cerebellum of 8-month-old MJD Tg-ATXN3-69Q mice. Interestingly, Nakano-Doi et al. demonstrated that induced ischemic stroke leads to an increase in VE-cadherin promoter activity and protein levels. They proposed that VE-cadherin may be involved in BBB repair following insult (50). Given that, it is plausible that upregulation in VE-cadherin protein levels may be a compensatory mechanism of TJ disruption in MJD Tg-ATXN3-69Q mice.

Notably, MSC administration increased occludin levels while reducing claudin-5 and VE-cadherin levels in the membrane, the subcellular locations where these proteins exert their function. Additionally, MSC treatment partially improved ZO-1 connection with the cytoskeleton, re-establishing its expression to levels comparable to those in the control group in this fraction.

In line with our results, MSC’s potential to rescue BBB function has also been described in PD. For instance, Chao *et al*. demonstrated that IV transplantation of MSC could prevent dopaminergic neuron death and recover BBB integrity in a preclinical model of PD induced by 1-methyl-4-phenyl-1,2,3,6-tetrahydropyridine. Indeed, MSC were able to restore claudin-1, claudin-5, and occludin levels in the substantia nigra compacta (51). Likewise, MSC-based therapies have also been shown to reduce vascular dysregulations in acute neurodegenerative disorders and in cerebrovascular conditions marked by BBB disruption, such as ischemic stroke (reviewed in (52, 53)), cerebral small-vessel disease (54), intracerebral hemorrhage (55–57), and traumatic brain injury (58, 59). Future studies should explore the mechanisms through which MSC promote vascular improvement in polyQ disorders, particularly through AJ/TJ recovery.

Remarkably, the effects of MSC on AJ/TJ proteins displayed a sex-specific pattern. Of note, a recent meta-analysis that evaluated the therapeutic effects of MSC in Huntington’s disease models revealed that MSC could prevent weight loss, but only in male mice (60). Importantly, MSC are recognized for their heightened sensitivity and responsiveness to their microenvironment. Hence, it is plausible that the underlying sex dimorphism in PolyQ disorders pathogenesis could be a significant factor influencing the MSC effect. Notably, the changes in vascular structure, cerebellar blood volume, and clearance were more evident in male than in female Tg-ATXN3-69Q mice. Additionally, female Tg mice displayed higher α-SMA expression than their male counterparts, which suggests that females may have stronger compensatory mechanisms, mitigating some functional dysregulations. Indeed, it is important to note that estrogen can have a protective effect in the neurovascular unit by regulating blood flow, BBB permeability, and inflammation (61). Our findings contribute to accumulating evidence supporting sex-specific variations in neurodegenerative disorders (review in (62)). Although research on this topic is currently limited, clinical evidence suggests that in PolyQ disorders, symptoms may progress faster in women than in men (63–67). Additionally, in a recent study, our group showed an increase in total cerebellar volume in MJD female patients. Contrarily, an increase in total cerebellar CSF volume was reported in MJD male patients, but not in females (68). Additionally, preclinical studies in HD mouse models demonstrate more severe HD dysregulations in males, which is in line with the observations in the present study (69–71). Such variations in disease progression may be attributed to natural sex dimorphisms in key cellular mechanisms that are relevant to neurodegenerative disorders, such as inflammation, circadian cycle, mitochondrial (dys)functioning, and BBB regulation (61, 70, 72, 73). In the future, investigations on PolyQ disorders, and specifically in the context of MJD, should also address potential sex differences and clarify the contradictions in current evidence.

On the other hand, while the influence of sex (of treated subjects) on the therapeutic efficacy of MSC remains largely unexplored in neurodegenerative disorders, existing studies have identified sex differences in MSC therapeutic potential in the context of non-neurological conditions, such as chronic inflammatory bone loss and osteoporosis (74, 75). Additionally, the influence of sex hormones, such as 17β-estradiol and testosterone, should also be considered as they could alter MSC secretion profile and consequently their protective function (76–78). Collectively, these findings emphasize the need for future investigations into the nuances of MSC effects across different demographic groups in MJD and other neurodegenerative disorders.

In conclusion, our study reveals that neurovascular dysfunction occurs in MJD models earlier than previously reported (9), characterized by significant morphological and functional vascular impairments, including BBB breakdown. We show that alterations in the expression and localization of key AJ/TJ-associated proteins in MJD mice precede and likely contribute to BBB leakage. Additionally, our findings suggest that MSC administration partially reverses vascular impairments in the Tg-ATXN3-69Q MJD model, demonstrating the multi-target therapeutic effects of MSC in addressing vascular disruptions. These insights not only enhance our understanding of MSC mechanisms in neurodegenerative disorders but also position MSC as a promising therapeutic strategy for mitigating neurovascular dysfunctions in MJD and potentially other PolyQ disorders.

## Abbreviations

BBB: Blood-brain barrier;
CoIV: Collagen type IV;
DAPI: 4′,6-diamidino-2-phenylindole;
EB: Evans blue;
GFAP: Glial fibrillary acidic protein;
HA: Hemagglutinin;
Iba1: Ionized calcium-binding adapter molecule 1;
MJD: Machado-Joseph disease;
MMP: Matrix metalloproteinase;
ROI: Region of interest;
SCA: Spinocerebellar ataxia;
SCA3: Spinocerebellar ataxia type 3;
TJ: Tight junction;
ZO-1: Zonula occludens-1

## Author contributions

IB: research study design, experiment conduction, data acquisition, data analysis, and manuscript writing; DG: data acquisition and data analysis; DL: data acquisition and data analysis; SL: data acquisition and data analysis; IM: data acquisition; AS: data acquisition; CH: data analysis; CG: research study design; RN: experiment conduction; L.P.d.A: research study design, provision of reagents, manuscript writing and reviewing; COM: research study design, experiment conduction and manuscript writing and reviewing. All authors contributed to the article and approved the submitted version.

## Consent for publication

All authors confrm their consent for publication

---

**Supplementary Figure 1.** MSC reached the cerebellum of Tg-ATXN3-69Q mice, despite vascular abnormalities.

Schematic representation of the experimental design (A): mice received an IV injection of MSC-Luc, then 10 min later received an IP injection of luciferin; bioluminescent signal was captured 30 min after MSC administration in an IVIS Lumina XR equipment. Representative bioluminescence image of MSC-luc signal in the brains of Tg-ATXN3-69Q mice and the evaluated region of interest (B). Values of bioluminescence (photons/s) in the brains of Wt T and Tg T mice (Wt T n=10 [9♂+1♀] vs Tg T n=11 [6♂+5♀]) (C). *Data presented as mean ± SD. unpaired t-test, p>0.05 = not significant*.

**Supplementary Figure 2.** Vascular dysregulations in the cerebella of 2.5-month-old Tg-ATXN3-69Q mice ICG signal intensity was detected in the cerebellar region of mice (Wt n=18 [9♂+9♀] vs Tg n=17 [11♂+6♀]) 10 min upon injection ICG at week 0 (A). Clearance of ICG after injection was evaluated by acquiring cerebellar ICG signal intensity assessed 10, 30, and 60 min in mice (Wt n=8 [4♂+4♀] vs Tg Nt n=6 [3♂+3♀] vs Tg n=8 [3♂+5♀]) upon injection at week 0 (B). Representative WB membranes and quantification of the expression levels of α-SMA (Wt n=9 [6♂+3♀] vs Tg n=13 [8♂+5♀]) (C) in the cerebellum 2.5-month-old mice. Protein relative levels were normalized with GAPDH. Representative images and quantification of the surface area of CoIV (Wt n=6 [3♂+3♀] vs Tg n=6 [3♂+3♀]) (D) and of vWF (Wt n=6 [4♂+2♀] vs Tg n=7 [5♂+2♀]) (E) staining in the cerebellum of 2.5-month-old. Data presented as mean ± SD. Unpaired t-test and two-way ANOVA followed by Tukey test. p>0.05 = not significant; * p<0.05, **p< 0.01, ***p< 0.001, ****p< 0.0001.

**Supplementary Figure 3.** BBB permeability is increased in 8-month-old Tg-ATXN3-69Q mice cerebella.

Representative WB membrane and quantification of Albumin protein levels in the cerebellum of 2.5-(Wt n=12 [9♂+3♀] vs Tg n=11 [7♂+4♀]) (A) and 8-month-old (Wt n=13 [6♂+7♀] vs Tg n=13 [8♂+5♀]) (B) mice. Protein relative levels were normalized with GAPDH. Representative images and quantification of the surface area of extravascular fibrinogen in the cerebellum of 2.5- (Wt n=5 [2♂+3♀] vs Tg n=7 [5♂+2♀]) (C) and 8-month-old (Wt n=4 [2♂+2♀] vs Tg n=6 [3♂+3♀]) (D) mice. Vascular fibrinogen is shown in white (co-localization of magenta and yellow) and CoIV in magenta, whereas extravascular fibrinogen is seen in yellow. Spearman’s correlation between CoIV and extravascular fibrinogen surface area coverage (Wt n=4 [2♂+2♀] vs Tg n=6 [3♂+3♀]) (E). Data presented as mean ± SD. Unpaired t-test. p>0.05 = not significant; * p<0.05.

**Supplementary Figure 4.** Longitudinal dysregulations of TJ-associated proteins expression in the cerebella of Tg-ATXN3-69Q mice

Representative WB membranes and quantification of the expression levels of TJ protein occludin 65 (A-F), Claudin-5 isoforms (G-O) of 27 (J-L) and 100 kDa (M-O), and the TJ cytoplasmatic adaptor ZO- 1 (P-U) in total protein extracts from 0.5- (Wt n=7 vs Tg n=7), 2.5- (Wt n=8 vs Tg n=7) and 8-month-old mice (Wt n=11-12 vs Tg n=11-12). Protein relative levels were normalized with GAPDH. Data presented as mean ± SD. Unpaired t-test. p>0.05 = not significant; **p< 0.01, ***p< 0.001, ****p< 0.0001.

**Supplementary Figure 5.** Longitudinal dysregulations of VE-cadherin expression in the cerebella of Tg-ATXN3-69Q mice

Representative WB membranes (A-C) and quantification of the expression levels of the AJ VE-cadherin (D-F) in total protein extracts from 0.5- (Wt n=7 vs. Tg n=7), 2.5- (Wt n=8 vs. Tg n=7) and 8-month-old mice (Wt n=12 vs. Tg n=13). Protein relative levels were normalized with GAPDH. Data presented as mean ± SD. Unpaired t-test. p>0.05 = not significant; * p<0.05.

**Supplementary Figure 6.** Correlation between TJ subcellular localization levels and motor as well as gait performance.

Pearson correlation between latency to cross the beam and: occludin 65kDa levels in membrane protein extracts (Wt n=11 vs Tg Nt n=4 vs Tg T n=5) (A), claudin-5 27kDa levels in membrane protein extracts (Wt n=9 vs Tg Nt n=4 vs Tg T n=5) (B), and ZO-1 levels in cytoskeleton protein extracts (Wt n=6 vs Tg Nt n=3 vs Tg T n=3) (C). Pearson correlation between footprint left foot overlap: occludin 65kDa levels in membrane protein extracts (Wt n=11 vs Tg Nt n=4 vs Tg T n=4) (D), claudin-5 27kDa levels in membrane protein extracts (Wt n=11 vs Tg Nt n=4 vs Tg T n=4) (E), and ZO-1 levels in cytoskeleton protein extracts (Wt n=6 vs Tg Nt n=3 vs Tg T n=2) (F). p>0.05 = not significant; **p< 0.01; ***p< 0.001; ****p< 0.0001.

**Supplementary Table 1.** Summary of AJ/TJ dysregulation in Tg-ATXN3-69Q mice discriminated by sex

## Supplemental methods and materials

### Behavioral Assessment

#### Footprint test

Gait was evaluated by performing footprint analysis, in which the paws of the animals were painted with red and blue non-toxic dyes on the hind and forefeet, respectively. Mice walked along a 100-cm-long, 10-cm-wide runway (with 15-cm-high walls). A fresh sheet of white paper was placed on the floor of the runway for each run. Five consecutive steps (excluding the footprints at the beginning and end of the run) were chosen for the analyses of the following parameters: stride length, the average distance of forward movement between each stride; overlap, the distance between the placement of and hind footprints on each side; hind-base with and front-base with, measured as the distance between left and right hind or forward footprints, respectively. The mean values of four measurements were used for statistical analysis.

#### Beam Walking test

Motor coordination and balance were evaluated using the beam walking test, where the mean latency time that mice spent crossing an 18 mm width square beam for 40 cm was recorded. The beam apparatus was 25 cm high with only one end attached to an escape box. Two trials were performed and the mean time they spent to cross the beam is represented.

### MSC-luciferase biodistribution evaluation

#### MSC infection with lentivirus encoding for Luciferase

MSC were infected with lentiviral vectors encoding for luciferase (pLenti PGK V5-LUC Puro plasmid construction, with puromycin resistance gene, Addgene), as previously reported in (Oliveira Miranda et al., 2018). Briefly, lentiviruses were produced in human embryonic kidney (HEK) 293T cells using a four-plasmid system as previously described (de Almeida et al., 2002). Lentiviral particles were resuspended in 0,5% bovine serum albumin (BSA) in phosphate-buffered saline solution (PBS) and viral particle content was determined by qPCR amplification. Viral stocks were stored at −80 °C until use.

MSC with 13 passages were plated in 6 multiwell plates (200,000 cells/well) and 24h later, cells were infected with 30 ng of virus/ 100,000 cells lentivirus encoding for luciferase, plus 4 μL/mL of hexadimethrine bromide (Sigma). MSC were incubated overnight at 37 °C in a 5% CO2/air atmosphere, and the culture medium was replaced to remove the lentivirus. MSC expressing luciferase (MSC-Luc) were selected by incubation with 0.01 mg/mL puromycin (Gibco), and then MSC-Luc were allowed to recover in a complete medium. Luciferase expression was confirmed by bioluminescence imaging performed using an IVIS Lumina XR (Caliper Life-Sciences, Hopkinton).

#### MSC biodistribution after intravenous injection

For the bioluminescence study, mice were treated with MSC-luc following the same experimental design. Images were acquired 30 min after mice (both wild-type and Tg) received an IV injection of 6 x 107 MSC-Luc/kg resuspended in HBSS in an IVIS Lumina XR imaging system using Living Image software (version 4.10; Xenogen). For each determination, mice were injected intraperitoneally with D-luciferin (100 mg/kg) and anesthetized with a non-lethal dose of a mixture of ketamine (Clorketam 1000, Vétoquinol, Lure, France) and xylazine (Rompun, Bayer, Leverkusen, Germany). Bioluminescence images were obtained 20 min after D-luciferin injection. For the quantification of the bioluminescent signal, regions of interest (ROI) were drawn around the cranium.

### Protein extraction

#### Protein extraction: Total Protein

Protein extracts were obtained from the right cerebellar hemisphere. For that, the tissue was first homogenized using Ambion TRIzol reagent (Fisher Scientific) and subjected to a density gradient with chloroform to remove RNA. Then 100% ethanol was added and samples were centrifuged to sediment DNA. Subsequently, the protein-rich phenol-ethanol phase was carefully collected and isopropanol was added to precipitate protein. After centrifugation, the resulting pellet was washed with guanidine-ethanol solution and 100% ethanol. The dried pellet was solubilized in urea/dithiothreitol (DTT) solution supplemented with protease inhibitors (Roche Diagnostics), followed by incubation at 95 °C for 3 min. Total protein extracts were stored at −80 °C until further analysis.

Total protein extracts were further submitted to 2 series of 4 s of ultrasound pulses (1 pulse/s). Protein concentrations were determined using Bradford protein assay (Bio-Rad). Samples were denatured with sample buffer (9.3% DTT, 10% sodium dodecyl sulfate (SDS), 30% glycerol in 0.5 M Tris-HCl/0.4% SDS - pH 6,8 and bromophenol blue 0.012%) and incubated at 95 °C for 5 min.

#### Protein extraction: Subcellular Protein Fractioning

Tissue fractionation was performed according to a standard Subcellular Protein Fractionation Kit for Tissues (87790, Thermo Scientific) that enables stepwise isolation of cytoplasmic, membrane, nuclear-soluble, nuclear chromatin-bound, and cytoskeletal protein extracts. Briefly, 23-27 mg of cerebellar tissue (average size of 2.5- and 8-month-old Tg mice cerebella, respectively) was homogenized in 500 µL of cytoplasmic extraction buffer containing protease inhibitors, filtered (250 µm strainer), and centrifuged (500 × g, 4 °C, 5 min). The resultant supernatant, the cytoplasmic fraction, was collected, and the pellet was resuspended in 325 µL of ice-cold membrane extraction buffer containing protease inhibitors. After incubation (4 °C, 10 min), samples were centrifuged (3000 × g, 5 min) and the supernatant, the membrane fraction, was collected. The pellet was then resuspended in 110 µL ice-cold nuclear extraction buffer containing protease inhibitors, incubated with gentle mixing (4 °C, 30 min), and centrifuged (4000 × g, 4 °C, 5 min). The supernatant containing the soluble nuclear protein fraction was collected and the resulting pellet was resuspended in 80 µL RT nuclear extraction buffer containing protease inhibitors, CaCl2, and Micrococcal Nuclease. After incubation (RT, 30 min), samples were centrifuged (12000 x g, 5 min), and the chromatin-bound nuclear protein fraction supernatant was collected. Lastly, the pellet was resuspended in 60 µL RT pellet extraction buffer, incubated (RT, 10 min), and centrifuged (12000 x g, 5 min). The final supernatant containing cytoskeletal extract was collected. All samples were stored at −80 °C until further analysis.

Extracts obtained with this kit were compatible with Western blot (WB) and were free of contamination with proteins of other fractions, confirmed assessing the presence or absence labeling specific protein markers of each subcellular location by WB (unpublish data) Protein concentrations were determined using Bradford protein assay (Bio-Rad). Samples were denatured with sample buffer (9.3% DTT, 10% sodium dodecyl sulfate (SDS), 30% glycerol in 0.5 M Tris-HCl/0.4% SDS - pH 6,8 and bromophenol blue 0.012%) and incubated at 95 °C for 5 min.

#### Quantification of extravascular fibrinogen

To assess blood vessel leakage, the presence of extravascular fibrinogen was measured in mice cerebella as previously described (Drouin-Ouellet et al., 2015, Duarte Lobo et al., 2020). Briefly, tile images of sections co-stained against CoIV and fibrinogen were acquired with Plan-Apochromat 20X/0.8 M27 objective in Zeiss Axio Imager Z2 microscope with red (561 nm) and green (488 nm) laser, respectively. Three to four sagittal sections per animal (inter-section distance: 180 μm) corresponding to a similar cerebellar region were analyzed. Using Image J software, masks from CoIV (magenta) and fibrin (yellow) were obtained. Upon merging images in Adobe Photoshop 2019 (Adobe Systems Incorporated), magenta and white (corresponding to magenta and yellow co-localization) were removed, leaving only extravascular fibrinogen staining in yellow. The percentage of surface area for extravascular fibrinogen was then determined using Image J software.

