## Supplemental Table 1 for "Mesenchymal Stromal Cell Therapeutic Potential to Restore Neurovascular Integrity in Machado-Joseph Disease"

**Supplementary Table 1.** Summary of AJ/TJ dysregulation in Tg-ATXN3-69Q mice discriminated by sex

| Age | Proteins | | Wt | | | Tg | | | Wt *vs* Tg | | |
|---|---|---|---|---|---|---|---|---|---|---|---|
| | | | Male | Female | ♂ *vs* ♀ p | Male | Female | ♂ *vs* ♀ p | All | Male | Female |
| *0.5 M* | Occludin | 45 kDa | 0.6439±0.2135 | 1.267±0.5964 | # | 0.8864±0.2008 | 0.6135±0.2181 | > 0.05 | = | = [#] | = |
| | | 65 kDa | 0.8311±0.1753 | 1.127±0.4786 | > 0.05 | 0.5457±0.1326 | 0.7322±0.2869 | # | =/↓ | =/↓ | = [#] |
| | Claudin | 27 kDa | 0.8075±0.4695 | 1.144±0.3379 | > 0.05 | 1.251±0.0685 | 1.317±0.1811 | > 0.05 | = | = | = |
| | | 100 kDa | 0.7677±0.5862 | 1.174±0.4933 | > 0.05 | 0.5823±0.5062 | 0.9267±0.1811 | > 0.05 | = | = | = |
| | ZO-1 | 220 kDa | 0.8786±0.2706 | 1.091±0.5762 | > 0.05 | 0.3472±0.2139 | 0.8237±0.3533 | 0.0964 | = | =/↓ | = |
| | VE-Cadherin | 120 kDa | 0.7273±0.2774 | 1.205±0.1399 | *0.0291 | 1.239±0.1684 | 1.164±0.4076 | > 0.05 | = | =/↑ | = |
| *2.5 M* | Occludin | 45 kDa | 0.8309±0.4060 | 1.169±0.3734 | > 0.05 | 0.8324±0.3376 | 1.868±0.5824 | # | = | = | = [#] |
| | | 65 kDa | 0.8996±0.0399 | 1.100±0.0738 | **0.0030 | 0.5282±0.1086 | 0.6039±0.0697 | # | ↓↓↓ | ↓↓ | ↓↓ [#] |
| | Claudin | 27 kDa | 0.9750±0.1388 | 1.025±0.1379 | > 0.05 | 1.245±0.0813 | 1.262±0.0412 | > 0.05 | ↑↑ | ↑↑ | ↑ [#] |
| | | 100 kDa | 0.9836±0.0687 | 1.016±0.1229 | > 0.05 | 1.073±0.1551 | 0.9486±0.2157 | # | = | = | = [#] |
| | ZO-1 | 220 kDa | 0.9355±0.1500 | 1.064±0.2106 | > 0.05 | 1.016±0.0790 | 1.053±0.5033 | # | = | = | = [#] |
| | VE-Cadherin | 120 kDa | --- | --- | --- | --- | --- | --- | --- | --- | --- |
| *8 M* | Occludin | 45 kDa | 0.8184±0.2263 | 1.061±0.2354 | > 0.05 | 0.5146±0.1428 | 0.6408±0.2929 | > 0.05 | ↓↓ | ↓↓ | ↓↓ |
| | | 65 kDa | 1.010±0.4784 | 1.353±0.2871 | > 0.05 | 0.3216±0.0950 | 0.3170±0.1772 | > 0.05 | ↓↓↓↓ | ↓↓ | ↓ |
| | Claudin | 27 kDa | 0.9524±0.3265 | 1.024±0.2398 | > 0.05 | 1.844±0.6331 | 1.879±0.6135 | > 0.05 | ↑↑↑↑ | ↑↑↑↑ | ↑↑↑↑ |
| | | 100 kDa | 0.9999±0.1641 | 1.000±0.0757 | > 0.05 | 0.8675±0.1386 | 0.9210±0.1765 | > 0.05 | =/↓ | =/↓ | = |
| | ZO-1 | 220 kDa | 1.003±0.4151 | 0.9967±0.2388 | > 0.05 | 0.5733±0.3128 | 0.6396±0.2721 | > 0.05 | ↓↓ | ↓↓ | =/↓ |
| | VE-Cadherin | 120 kDa | 0.9880±0.2033 | 1.163±0.2528 | > 0.05 | 1.304±0.3117 | 1.463±0.4366 | > 0.05 | ↑↑ | =/↑ | =/↑ |

Statistical differences between male and female animals regarding AT/TJ levels in total protein cerebellar extracts of in 0.5-, 2.5- and 8-month-old mice of the Tg-ATXN3-69Q model and sex-specific patterns of dysregulation in Tg mice compared to Wt littermates (Wt *vs* Tg column).

↑↑↑↑ / ↓↓↓↓ represents an increase/decrease of more than 75%; ↑↑↑ / ↓↓↓ represents an increase/decrease of 50-75%; ↑↑ / ↓↓ represents an increase/decrease of 25-50%; ↑ / ↓ represents an increase/decrease of less than 25%; = represents an absence of changes; =/↓ represents a tendency to decrease without statistical significance. # - n too small to perform statistical analizes; ** $p < 0.01$. List of abbreviation: M – months; Tg – transgenic; Wt -wild-type; ZO-1 - zonula occludens-1; ♂ - male; ♀ - female.
